# Full-coverage native RNA sequencing of HIV-1 viruses

**DOI:** 10.1101/845610

**Authors:** Alejandro R. Gener, Jason T. Kimata

## Abstract

**Objective:** To evaluate native RNA sequencing for sequencing HIV-1 viral genomes

**Methods:** Fifteen HIV-1 strains were processed with Direct RNA Sequencing (SQK-RNA002) library kits and sequenced on MinION Mk1B devices with RevD flow cells (Oxford Nanopore Technologies (ONT), Oxford, UK). Raw reads were converted to FASTQ, aligned to reference sequences, and assembled into contigs. Multi-sequence alignments of the contigs were generated and used for cladistics analysis.

**Results:** We sequenced full-length HIV-1 from the transcriptional start site to 3’ LTR (100% virion genome) in 3 out of 15 isolates (89.6, NLAD8, AD17), achieving majority coverage (defined as > 50%) in another 7 out of 15 isolates. Inspection of NLAD8 sequence alignments revealed splicing or deletion signatures. Despite the strong 3’ bias, read coverage was sufficient to evaluate single-nucleotide variants (SNVs), insertions and deletions in 9 isolates, and to assemble HIV-1 genomes directly from viral RNA, achieving a maximum of 94% assembly coverage for NLAD8. Phylogenetic relationships were maintained at the level of contigs, as well as individual reads.

**Conclusions:** ONT native RNA sequencing performed as expected, covering full-length HIV-1 RNA without PCR or cDNA sequencing. Native single-molecule RNA sequencing supported previous models of HIV-1 replication, and samples exhibited strain-specific transcriptional signals. We propose Context Dependency Variant Classification to describe variants occurring in information-dense regions of HIV. These data provide rich resources for emerging RNA modification detection schemes. Future work will expand HIV-1 transcript profiling to infection models and clinical samples.

## INTRODUCTION

Human immunodeficiency virus type 1 (HIV-1) strains have high sequence variability due to their highly error-prone replication. Lack of full-length sequencing data has limited our understanding of HIV biology. The closest we have come to being able to observe the information contained in the HIV-1 viral genome directly has been to stitch together short read data into quasispecies [1], which are neither real nor direct observations of these viruses. Most approaches to sequencing the HIV-1 viral genome have used some variant of reverse transcription to make double-stranded cDNA, usually followed by PCR amplification, and finally cDNA sequencing (classic RNA-seq) [2]. Recently, we used long read PCR + DNA sequencing of synthetic plasmid-bound provirus to observe the entire HIV-1 DNA genome for the first time [3]. In addition to covering the HIV-1 reference sequence HXB2 deeply and evenly, and recovering at least one full-length read, we were able to observe 20 SNVs, half of which were in repetitive Long Terminal Repeat (LTR) regions. However, DNA-based sequencing methods cannot differentiate between reads from infectious virion RNA, integrated proviral DNA, and non-integrated forms [4]. Direct sequencing of the entire HIV-1 viral RNA genome with one full-length read of native virion-associated RNA has yet to be described. A barrier to full-coverage HIV-1 reads has been limitations in lengths of extracted nucleic acids from samples (Gener, unpublished). Additionally, HIV-1 exhibits complicated splicing behavior which may be virus-[5] and host cell-dependent [6]. The observation of alternative splicing of viral mRNAs [5] is coverage-dependent, often forcing investigators into using PCR to amplify signal at the expense of per-base accuracy. PCR also biases the signal that can be observed. The Oxford Nanopore Technologies (ONT) nanopore sequencers are the only commercially available platforms currently able to directly sequence RNA in its native form without the need for cDNA sequencing or PCR [7], and have been used to sequence other important RNA viruses like influenza A virus [8]. Here, we used MinION Mk1B devices to sequence 15 strains of full-length infectious HIV-1, with the goal of evaluating the technology for future applications.

## METHODS

### HIV-1 viruses

To generate infectious virus from plasmid clones of HIV-1 NL4-3, NLAD8, RHPA, and AD17, we transiently transfected 293T cells using ExtremeGENE 9 according to the manufacturer’s protocol (Roche). After 48 hours, supernatants were collected from transfected cultures of cells, passed through 0.45 μm-pore-size syringe filters, aliquoted, and saved at −80 °C. The amount of infectious virus was determined by limiting dilution on TZMbl cells using a luciferase reporter assay as we have described [9].

Other HIV-1 strains used in this study were derived from infectious virus obtained from the NIH AIDS Reagent and Reference Program (89.6, SF162, 97USNG30, MN, ADA, 92UG038, 92UG029, ELI, JR-FL, and BaL) or were generously provided by M.R. Ferguson and W.A. O’Brien (HIV1-SX) of the University of Texas Medical Branch, Galveston, TX. In short, peripheral blood mononuclear cells (PBMCs) were isolated from leukopacks of anonymous blood donors purchased from the Gulf Coast Blood Center (Houston, TX) using Lymphocyte Separation Medium (Sigma-Aldrich). Cultures of PBMCs were stimulated with phytohemagglutinin PHA-P and IL-2 (50 U/ml) in RPMI1640 complete (supplemented with 10% heat-inactivated fetal bovine serum, 2 mM L-glutamine, and 100 U/ml penicillin/ 100 μg/ml streptomycin) for 2 days prior to inoculation with HIV-1. After infection, PBMCs were cultured for an additional 10-15 days in RPMI complete with 50 U/ml IL-2. Virus production was confirmed by testing supernatants for HIV-1 p24 by ELISA (Advanced Bioscience Laboratories). Supernatants were collected from infected cultures of PBMCs, passed through 0.45 μm-pore-size syringe filters, aliquoted and saved at −80 °C. The amount of infectious virus was determined by limiting dilution titration of the virus stocks on PHA-stimulated PBMCs and assay for infectious virus production after 10-14 days of infection by HIV-1 p24 ELISA.

### Viral RNA isolation

RNA was extracted using the MagMAX-96 viral RNA isolation kit (Applied Biosystems) with a Kingfisher Flex Purification System (ThermoFisher) described by the manufacturer with the following modifications: 200 μl of viral lysate was added to 200 μl of freshly prepared Lysis/Binding Solution and allowed to incubate for at least 15 minutes (no more than 1 hour). Each lysate sample was loaded into a well of a 96-well deep well Kingfisher plate and processed following kit instructions. Each sample was eluted into 50 ul of Elution Buffer in duplicate into individual DNA LoBind 1.5 ml microcentrifuge tubes (Eppendorf) for storage at 80 °C until use in sequencing.

### Long-read native RNA sequencing

Extracted RNA was sequenced across three MinION Mk1B sequencers (Oxford Nanopore Technologies (ONT), Oxford, UK). Briefly, 9 ul of extracted viral RNA eluate was carried over and processed with Direct RNA kit SQK-RNA002 following ONT protocol DRS_9080_v2_revF_22Nov2018, Last updated: 19/07/2019. This protocol recommended 1 round of cDNA “first-strand” synthesis for the purpose of breaking up RNA 2° structure, however these cDNA strands are ultimately not sequenced. The reverse transcriptase kit used: SuperScript IV First Strand Synthesis System (Invitrogen). The ligase used: T4 DNA ligase (NEB). Libraries were loaded onto one of 5 MinION RevD flow cells, pore version R9.4.1. and sequencing runs approximating 6 to 11 hours were completed with MinKNOW (version 3.5.5) on three laptop computers.

Runs were stopped manually in MinKNOW based on empirical assessment of HIV-1 sequence coverage while Live Basecalling (ONT) was enabled. Between runs, flow cells were washed with Flow Cell Wash Kit (EXP-WSH003) (Version: WFC_9088_v1_revB_18Sep2019). Flow cells exhibiting increasing failed reads, or abrupt changes in membrane potential (bottoming out at −250 mV) were replaced with new flow cells. Flow cell resting potential was set at −180 mV at the beginning of each run, and MinKNOW was used to adjust voltage automatically at mux scans every 30 minutes during the runs.

Raw data was basecalled (converted from FAST5 to FASTQ format) with Guppy high-accuracy model version 3.3.0, on a compute cluster. Mapped basecalled reads were fed into Canu version 1.8 implemented on a cluster [10]. The parameter genomeSize in Canu was set to be 1 Kb, and stopOnLowCoverage was set to 10 reads. SAMTOOLS Flagstat [11] and QualiMap [12] were used for mapping stats from BAM files from minimap2 [13] implemented in Galaxy (usegalaxy.eu) [14]. SnapGene version 5.0.4 was used to manually annotate contigs from Canu. Select regions were interrogated with multisequence alignments with MAFFT [15] and visualized with Archaeopteryx 0.9921 beta (170712).

After basecalling, read length and mean quality score are reported in a sequencing_symmary.txt file in MinKNOW. For skipped reads (reads not basecalled by timed or forced end of sequencing runs), basecalling in standalone Guppy provides the same sequencing_symmary.txt information. For HIV-1-mapping reads, this information is provided in **Figure 1**. To pull this information, unique read IDs aligning to AF033819 (with minimap2) were concatenated into a master list and used as follows: egrep -f MASTERLIST.txt sequencing_symmary.txt > HIV-mapping_sequencing_symmary.txt. Data was wrangled in Microsoft Excel for Mac version 16.29.1. Graphing was done with GraphPad Prism 8 for macOS version 8.3.0. As samples were not compared directly to each other, additional statistics were not calculated.

**Figure 1A:**
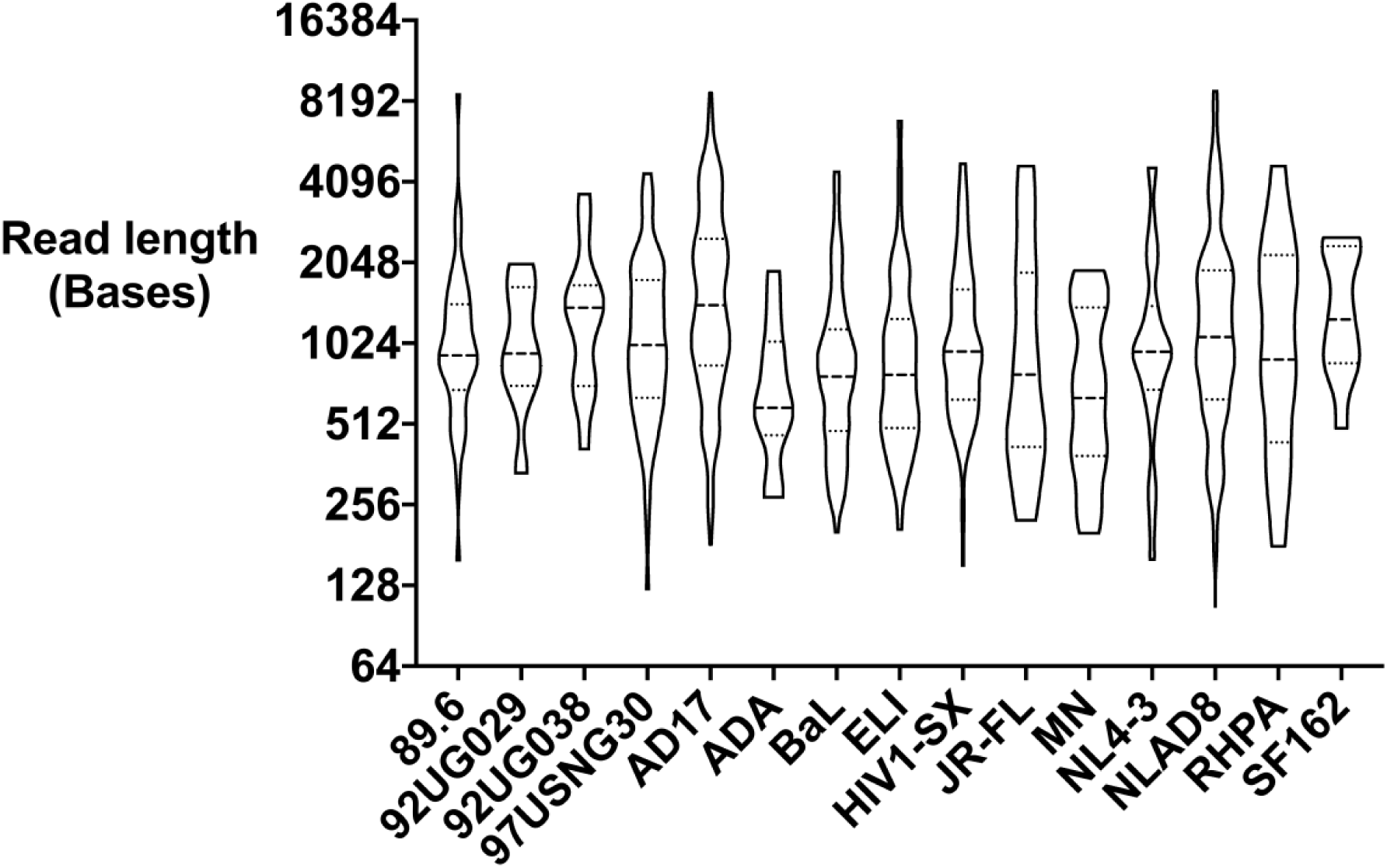
Read lengths for HIV-1-mapping reads sequenced in this study. Violin plot of read length. Log_2_-scaled. Median (big dash) and quartiles (little dash). All native RNA reads are longer than most short DNA-seq reads. Note: Reads from multiple runs either live basecalled or skipped reads basecalled afterward were collapsed. Per-run info is summarized in **Supplemental Table 1**.

**Figure 1B:**
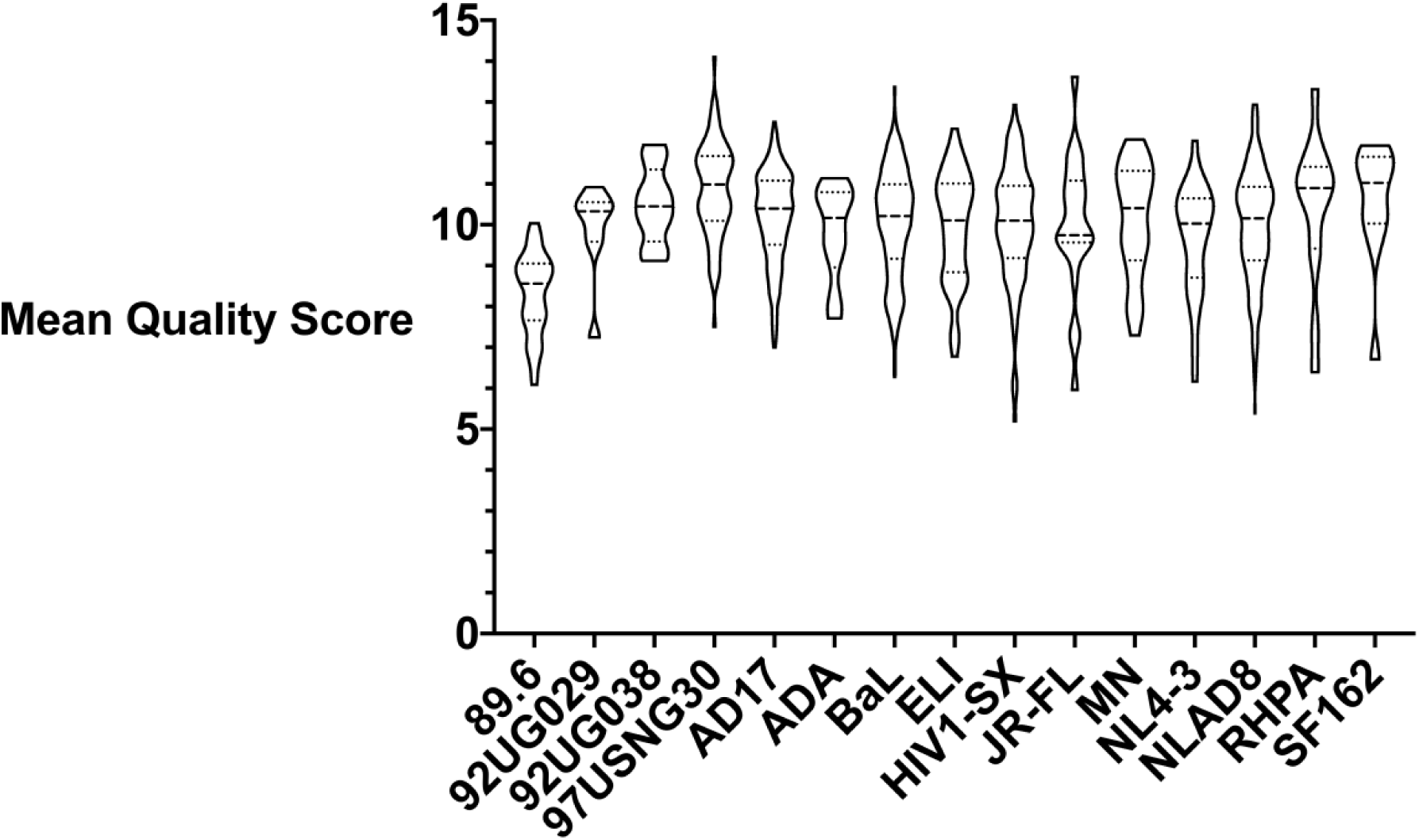
Read mean quality scores for HIV-1-mapping reads sequenced in this study. Violin plot of quality score. Median (big dash) and quartiles (little dash). Note: Reads from multiple runs either live basecalled or skipped reads basecalled afterward were collapsed. Per-run info is summarized in **Supplemental Table 1**.

### Reference sequences

The reference sequence of HIV-1 used in **Figure 2A-O** was from NCBI, accession number AF033819.3. **Supplemental Figure 11** used HXB2 K03455.1. See **Table 1** for additional reference accession numbers. The control RNA in SQK-RNA002 is ENO2, and the reference used for mapping was NM_001179305, a partial mRNA from Saccharomyces cerevisiae S288C phosphopyruvate hydratase ENO2 (ENO2), 1314 bp long.

**Figure 2A:**
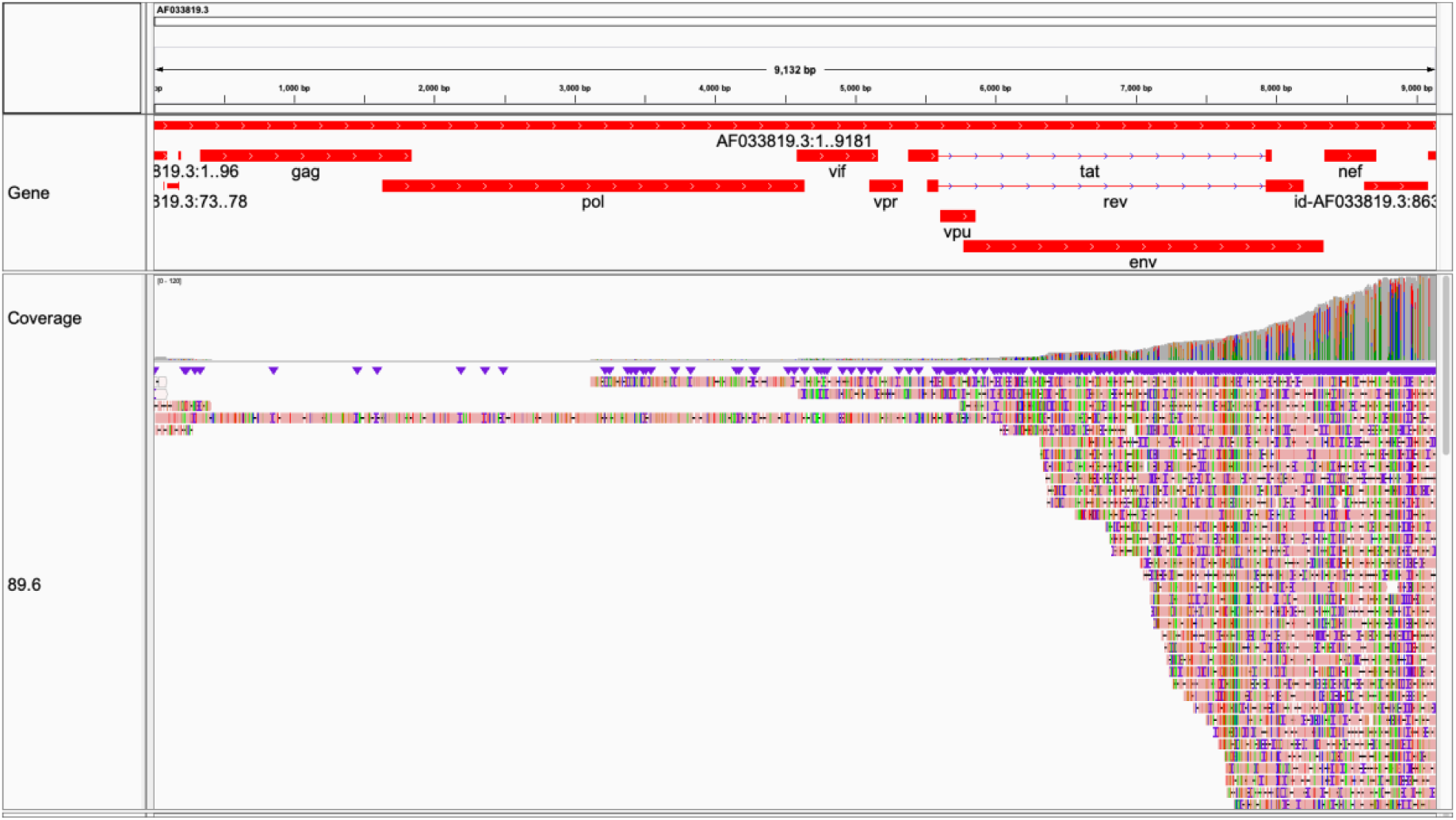
89.6. Full-coverage (9,166/9,181∼99.8%) achieved. Pink denotes forward strand orientation. Colors (Red, Green, Gold, Blue) in reads denote differences from reference, not necessarily read mismatches. Purple and black denote insertions and deletions in the reads, which may or may not be errors in individual reads. Mapping to reference was used to evaluate coverage across multiple HIV-1 strains. Note slight change in slope over RRE. Visualized in Integrative Genomics Viewer [49]. Coverage over *env* possibly sufficient for cladistics (Figure 4).

**Figure 2B:**
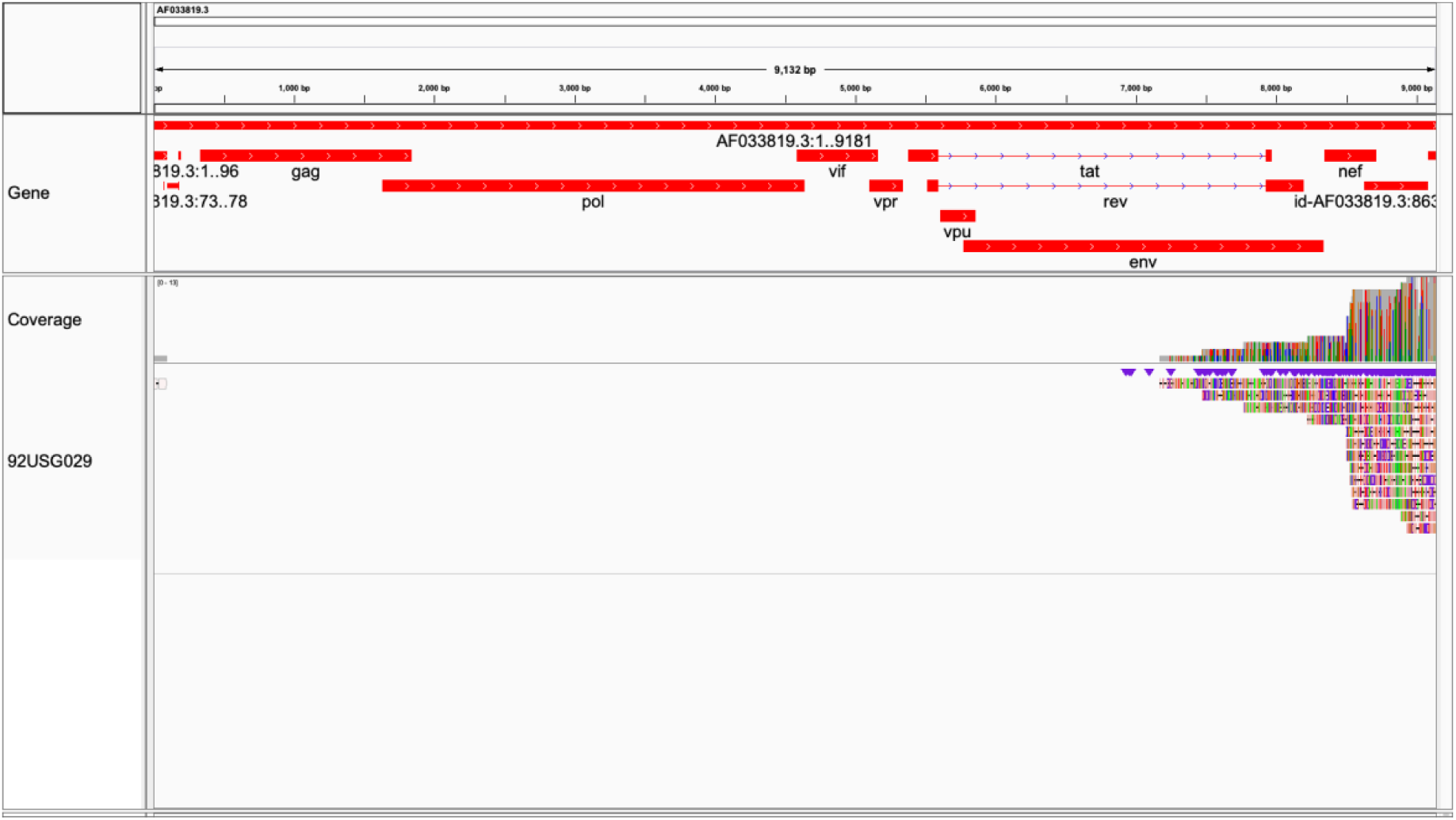
92UG029. Lower coverage (2,007/9,181∼21.9%) achieved. Visualized in Integrative Genomics Viewer [49]. Coverage over *env* insufficient for cladistics.

**Figure 2C:**
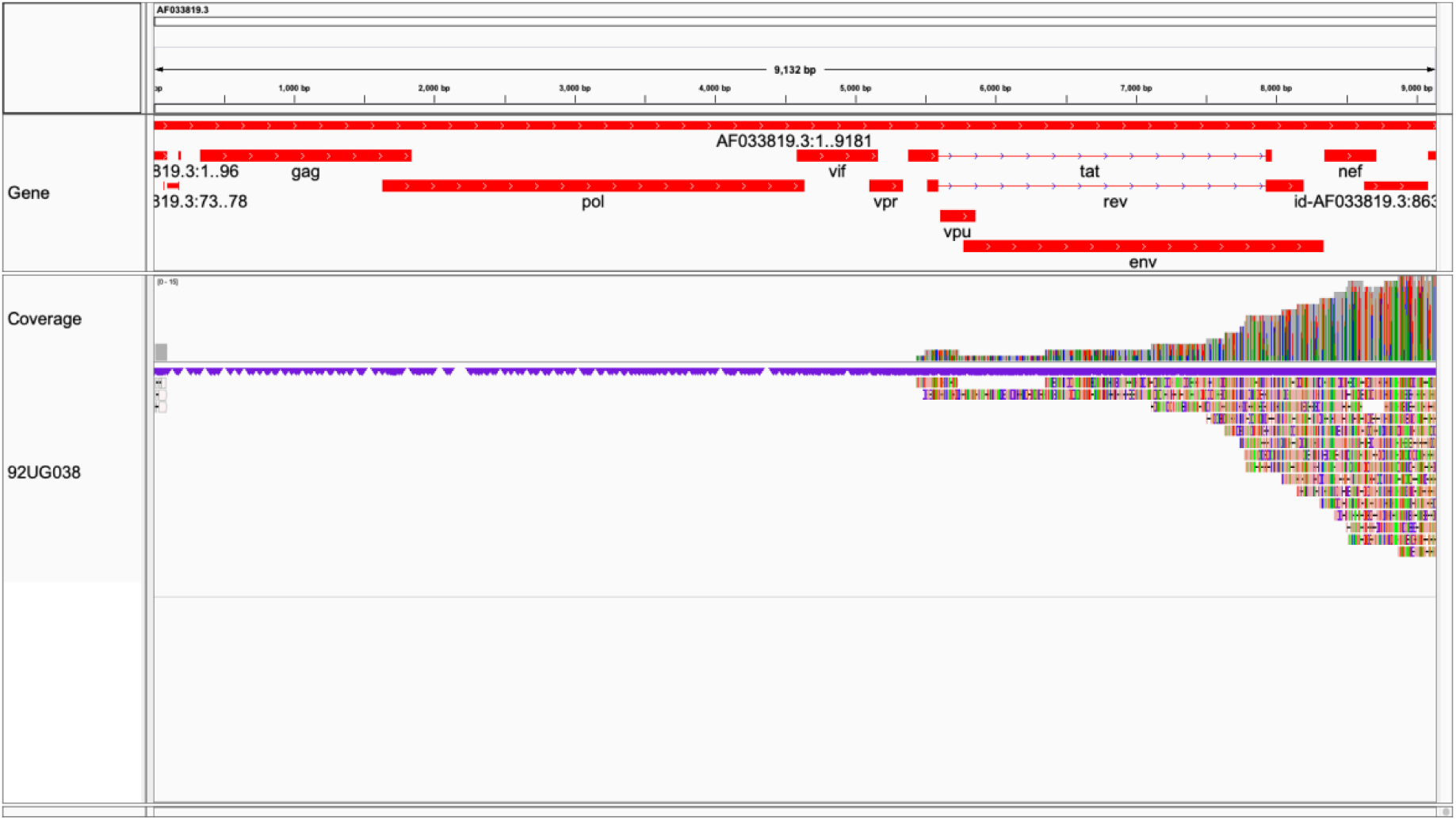
92UG038. Moderate coverage (3,678/9,181∼40.1%) achieved. Note slight change in slope over RRE. Visualized in Integrative Genomics Viewer [49]. Coverage over *env* insufficient for cladistics.

**Figure 2D:**
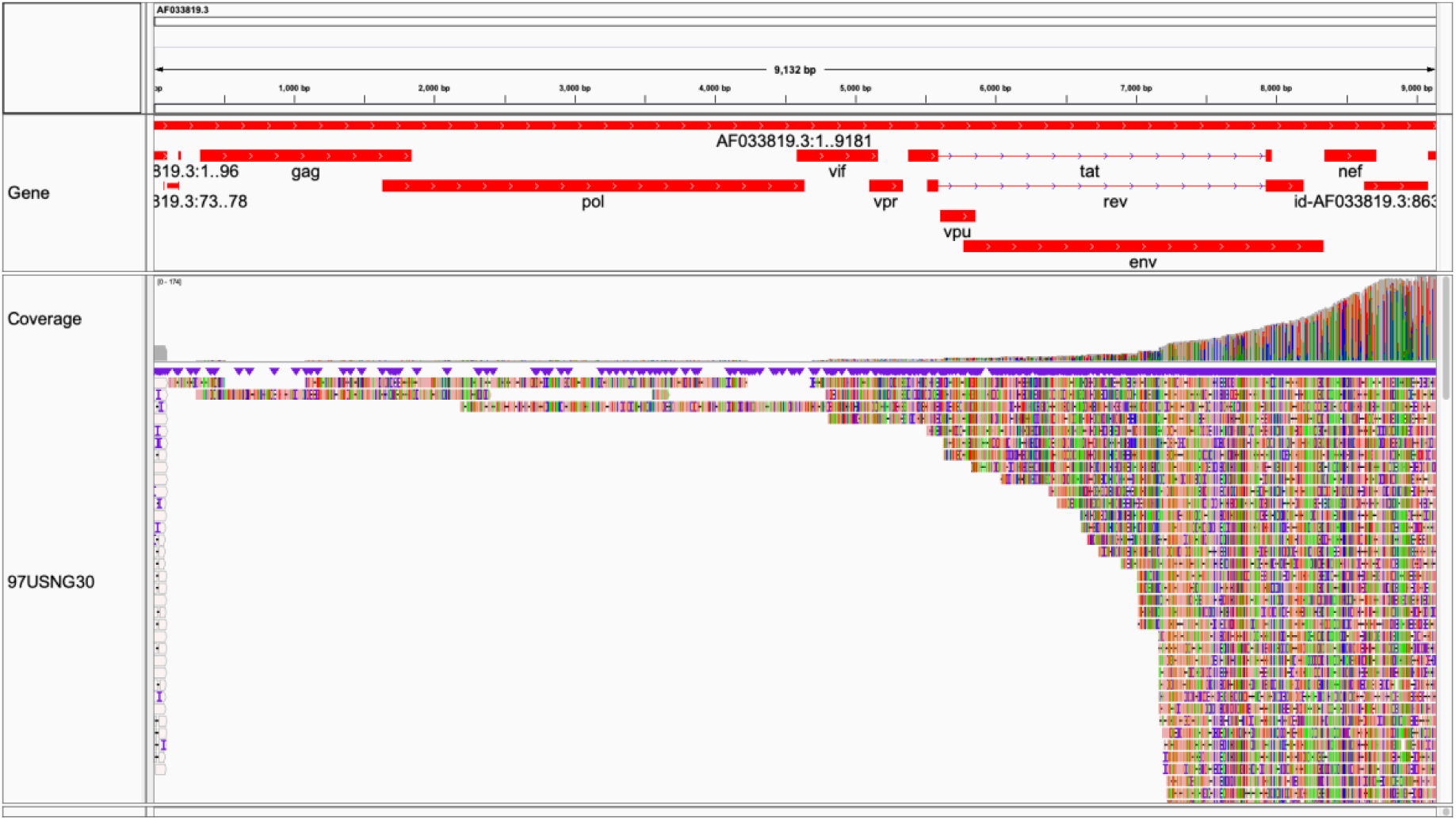
97USNG30. Majority coverage (6,992/9,181∼76.2%) achieved. Note slight change in slope over RRE. Visualized in Integrative Genomics Viewer [49]. Coverage over *env* sufficient for cladistics (Figure 4).

**Figure 2E:**
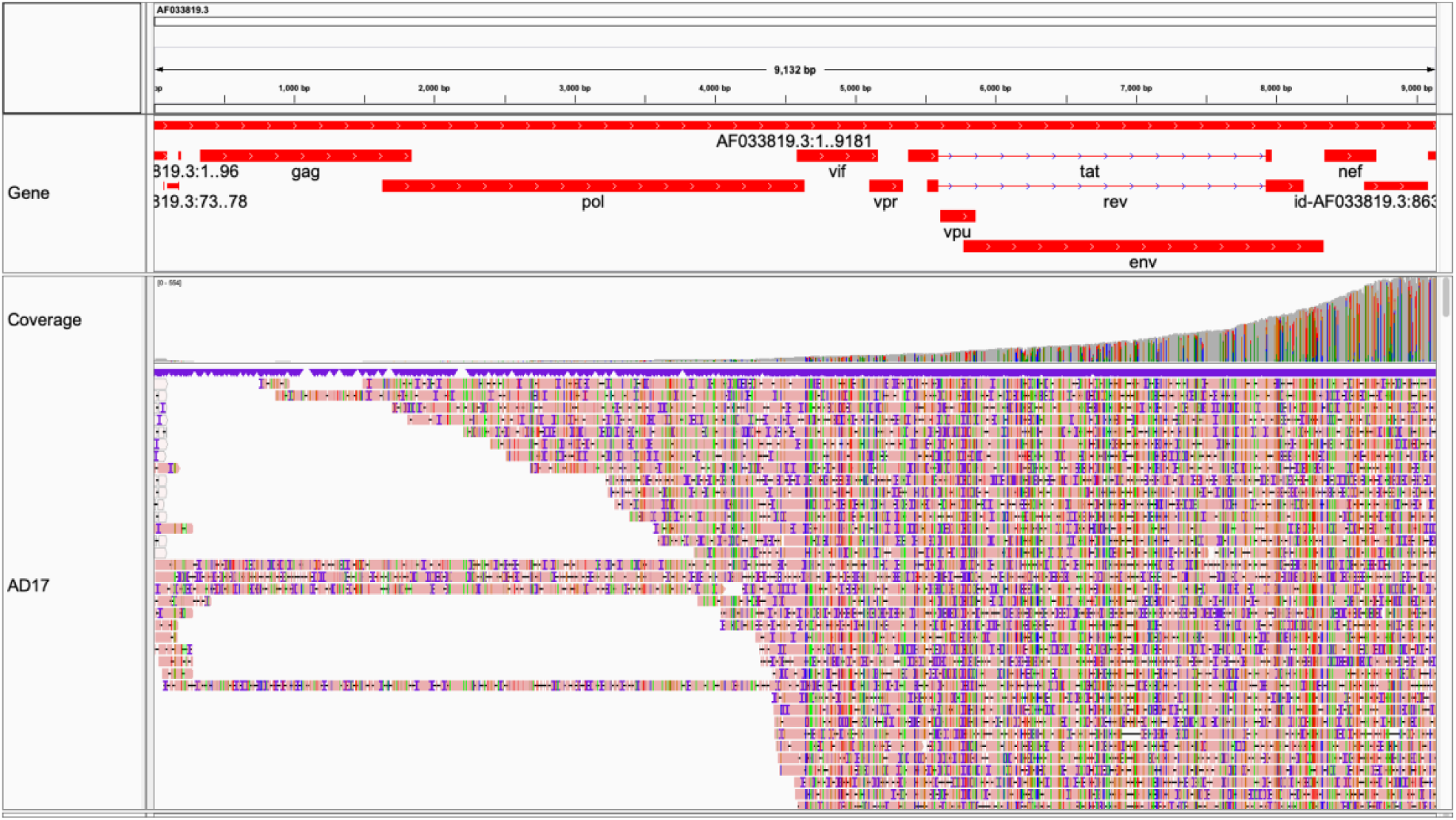
AD17. Full-length coverage (9,164/9,181∼99.8%) achieved. Visualized in Integrative Genomics Viewer [49]. Coverage over *env* sufficient for cladistics (Figure 4).

**Figure 2F:**
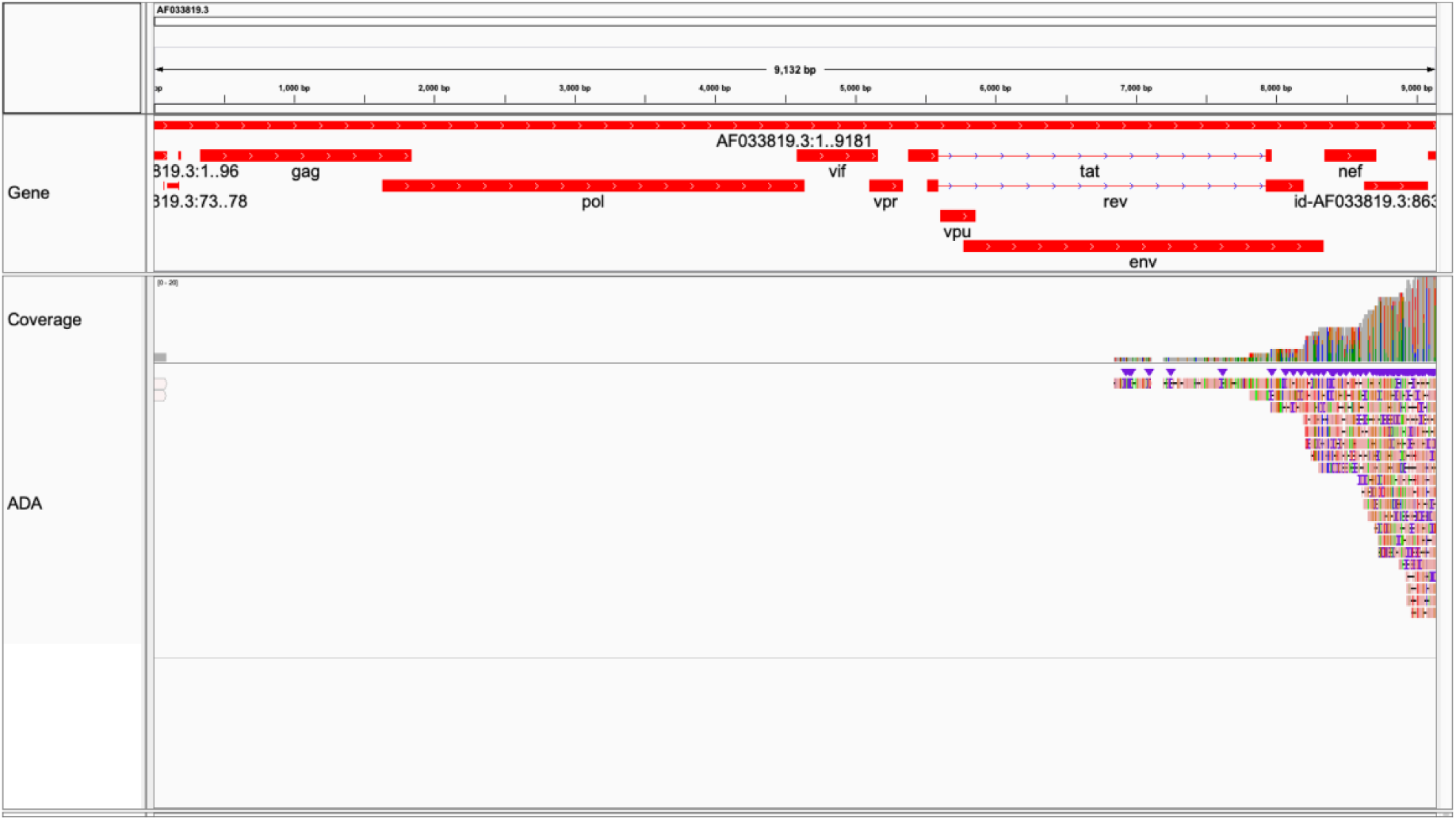
ADA. Lower coverage (1,981/9,181∼21.6%) achieved. Visualized in Integrative Genomics Viewer [49]. Coverage over *env* insufficient for cladistics (Figure 4).

**Figure 2G:**
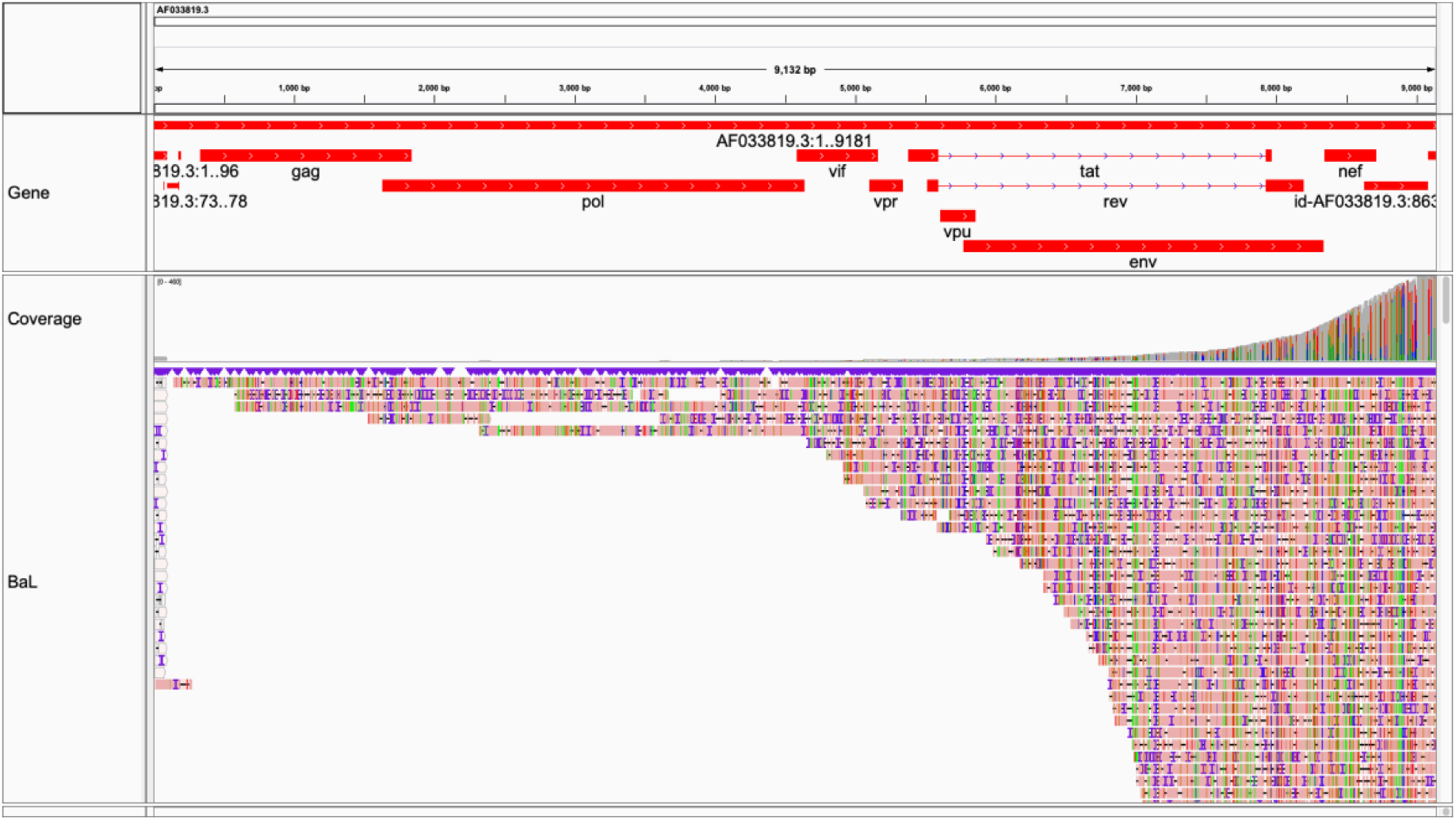
BaL. Majority coverage (8,597/9,181∼93.6%) achieved. Visualized in Integrative Genomics Viewer [49]. Coverage over *env* sufficient for cladistics (Figure 4).

**Figure 2H:**
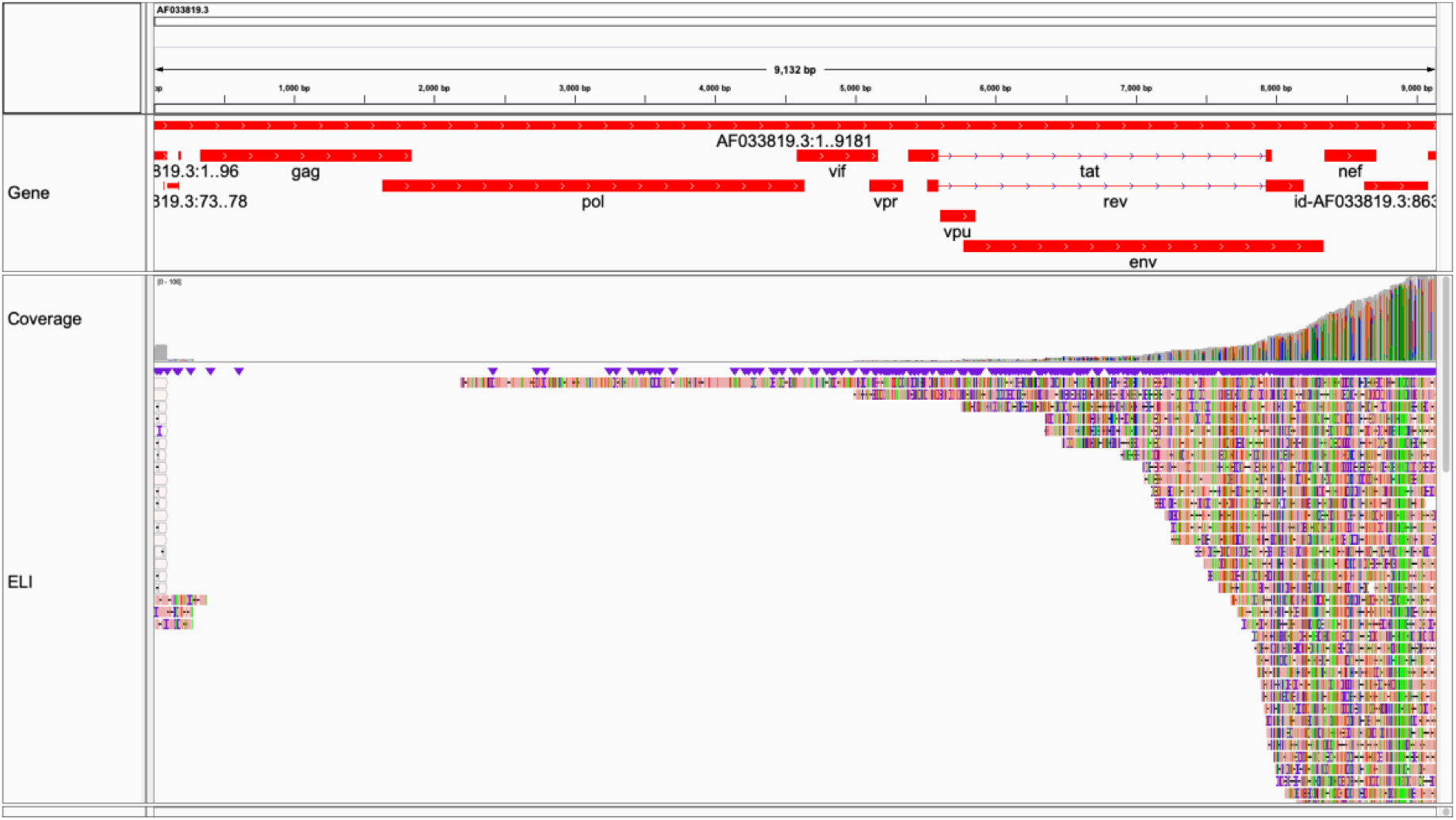
ELI. Majority coverage (6,987/9,181∼76.1%) achieved. Note slight change in slope over RRE. Visualized in Integrative Genomics Viewer [49]. Coverage over *env* possibly sufficient for cladistics (Figure 4).

**Figure 2I:**
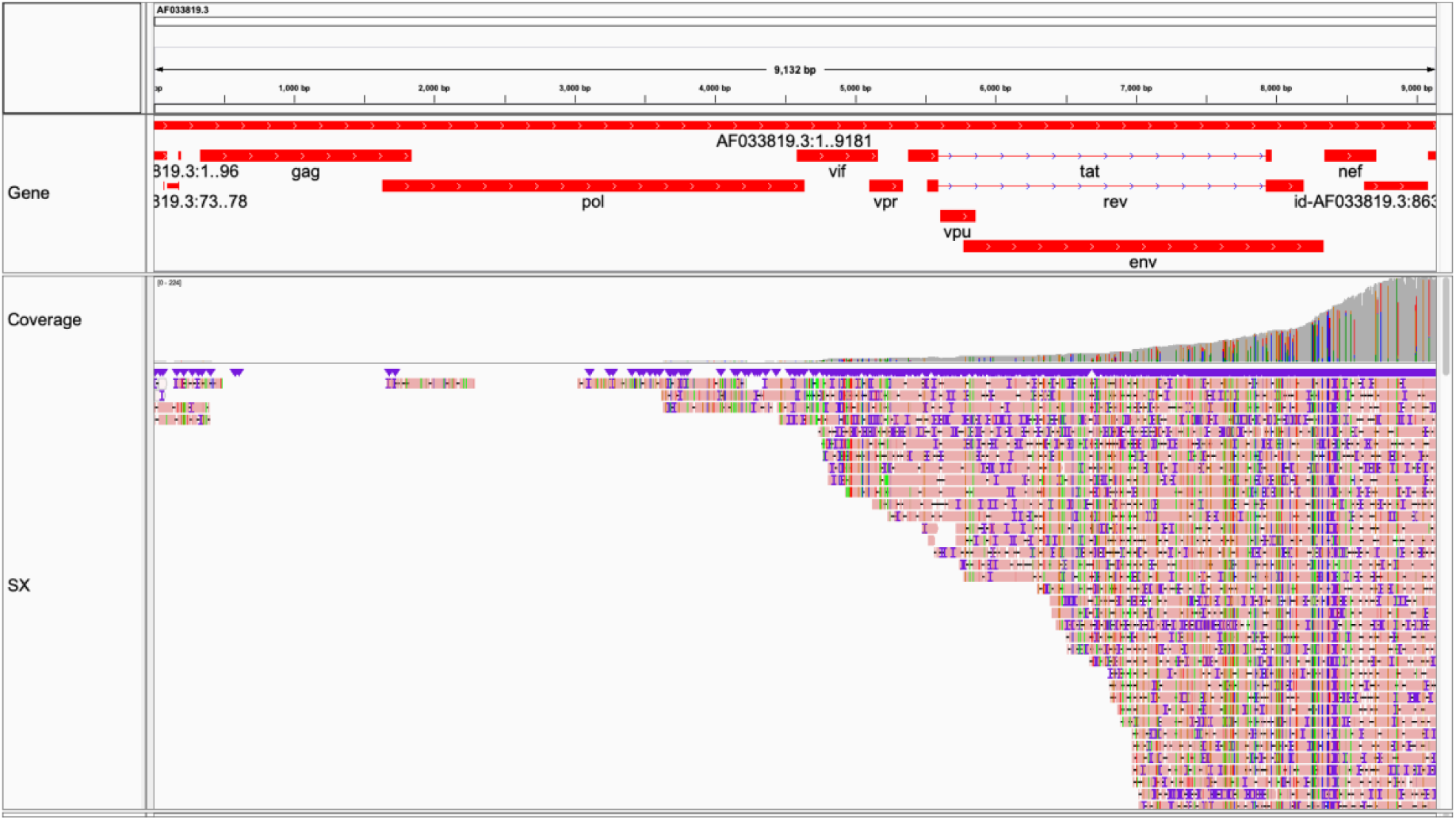
HIV1-SX. Majority coverage (5,561/9,181∼60.6%) achieved. Compare HIV1-SX, NLAD8, pNL4-3. Visualized in Integrative Genomics Viewer [49]. Coverage over *env* sufficient for cladistics (Figure 4).

**Figure 2J:**
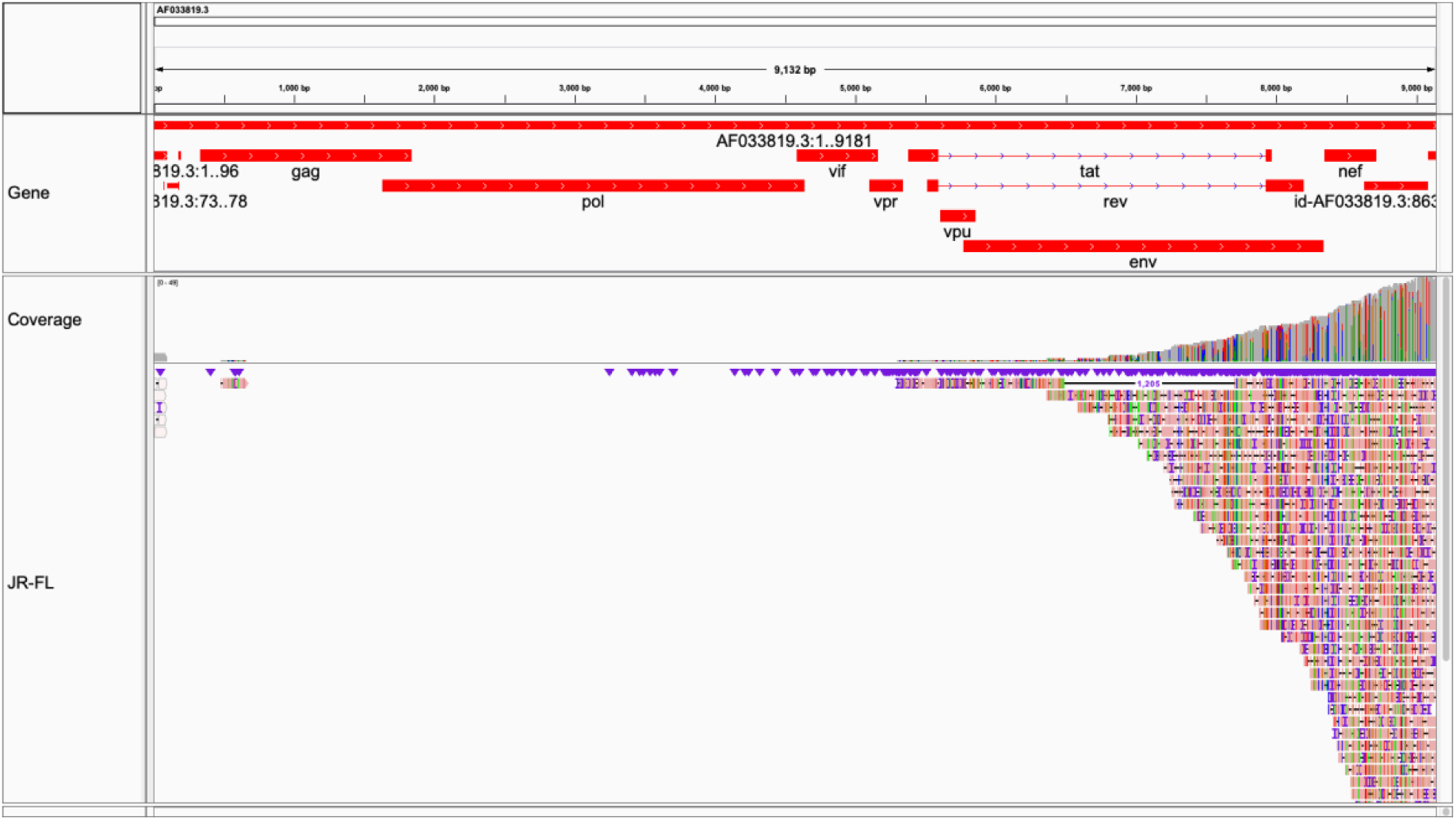
JR-FL. Moderate coverage (2,814/9,181∼30.7%) achieved. Visualized in Integrative Genomics Viewer [49]. Coverage over *env* insufficient for cladistics.

**Figure 2K:**
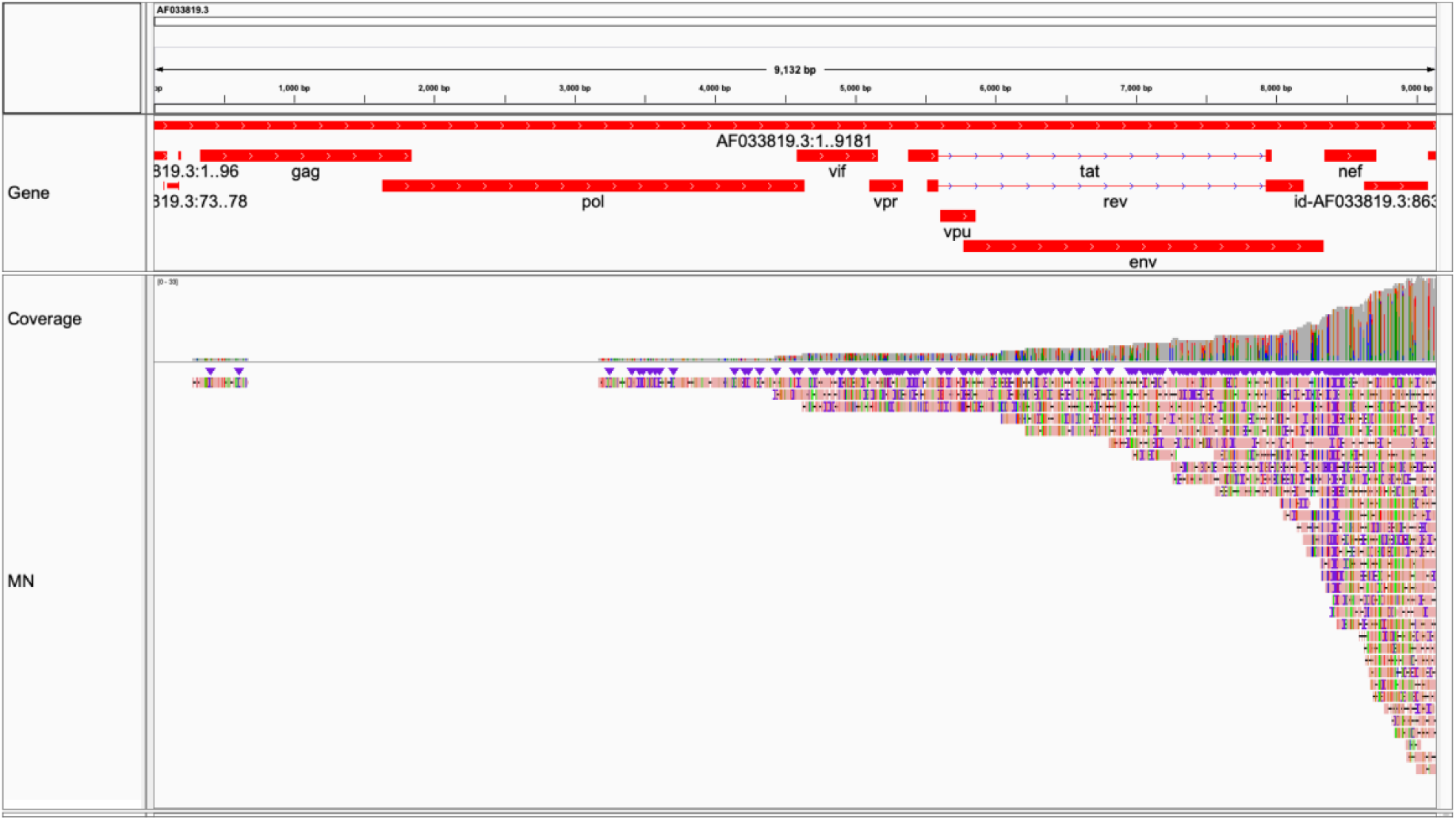
MN. Majority coverage (5,995/9,181∼65.3%) achieved. Visualized in Integrative Genomics Viewer [49]. Coverage over *env* possibly sufficient for cladistics (Figure 4).

**Figure 2L:**
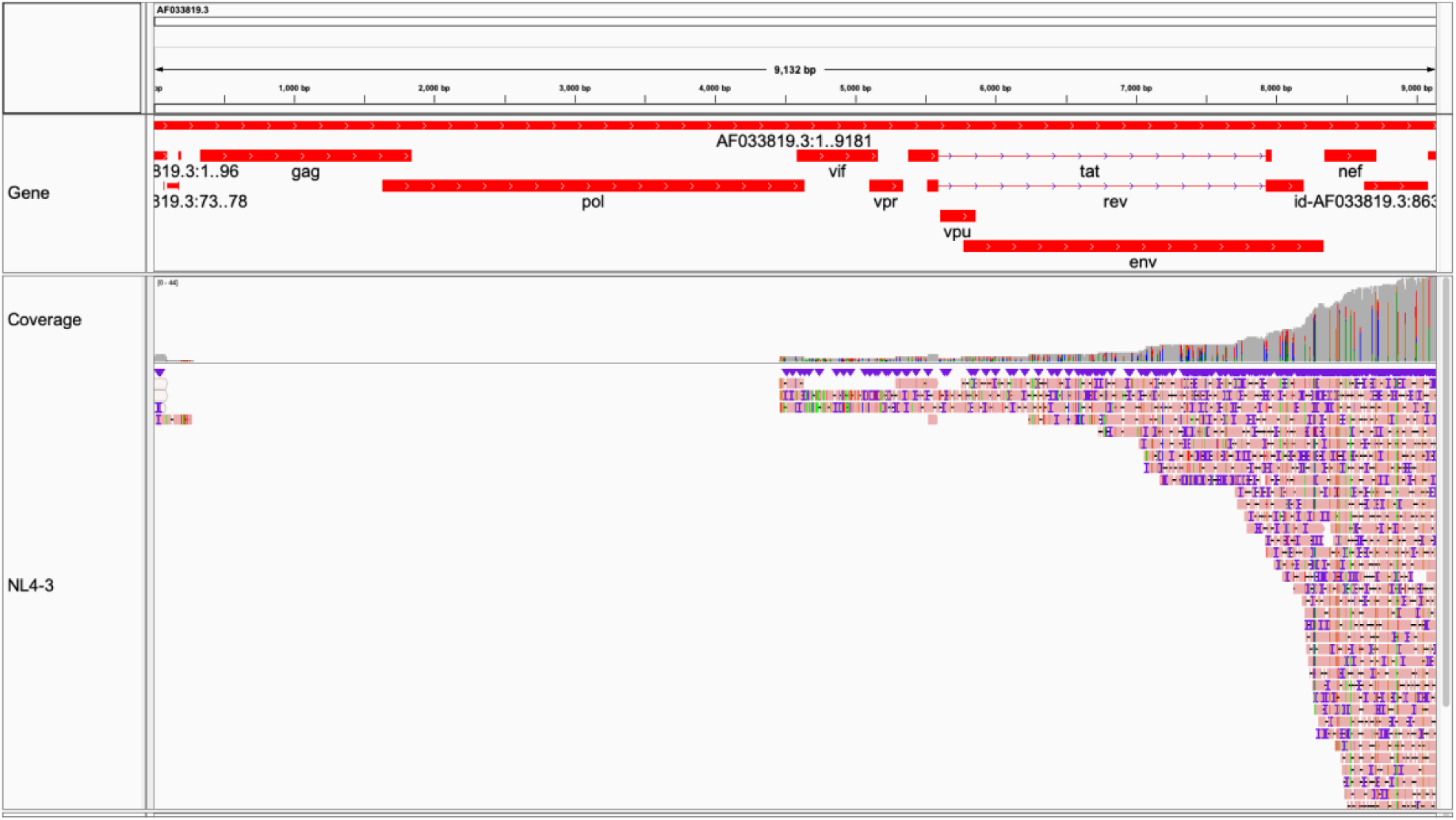
NL4-3. Majority coverage (4,715/9,181∼51.4%) achieved. Note slight change in slope over RRE. Visualized in Integrative Genomics Viewer [49]. Coverage over *env* possibly sufficient for cladistics (Figure 4).

**Figure 2M:**
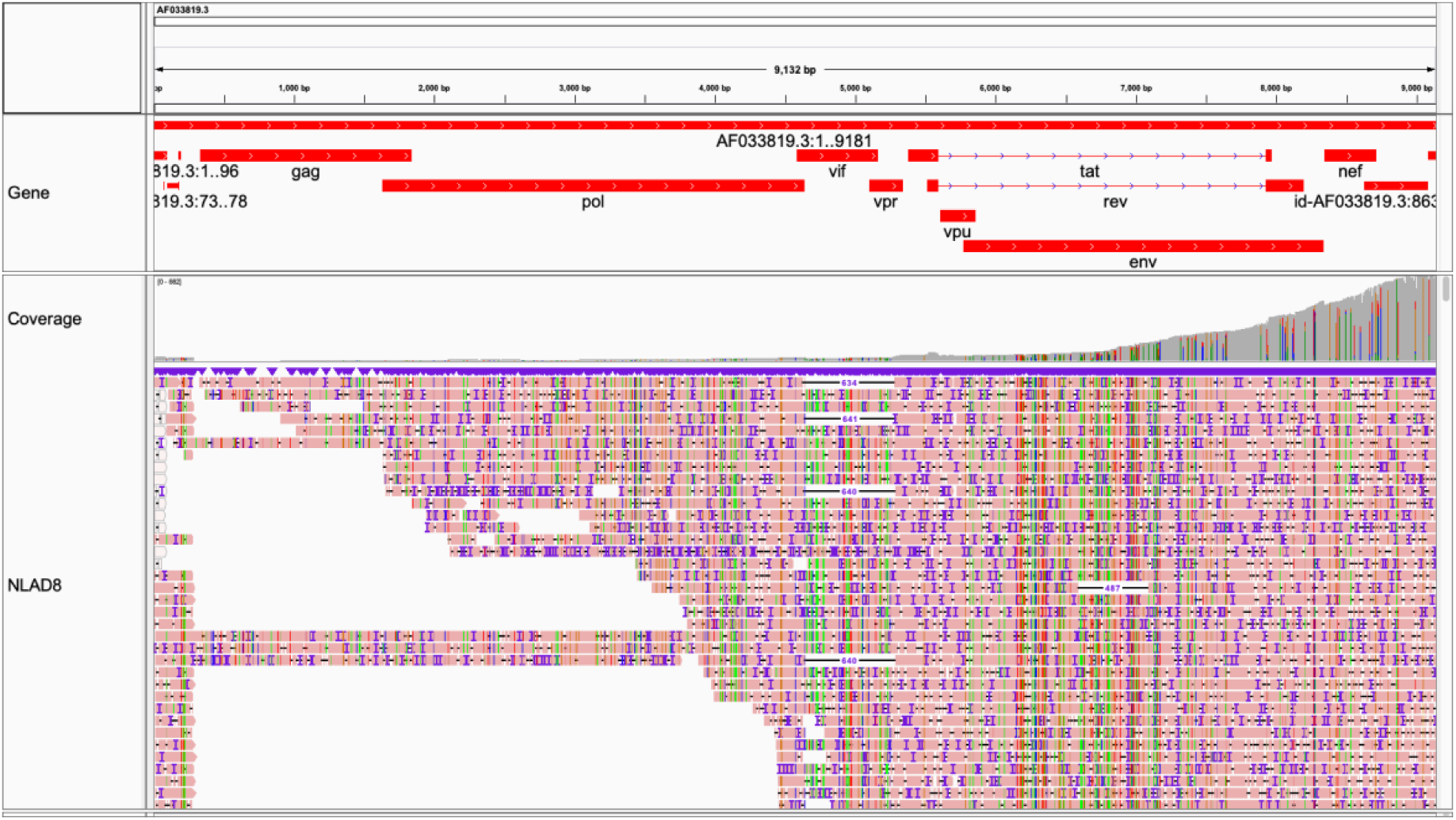
NLAD8. Full-length coverage (9,166/9,181∼99.8%) achieved. Visualized in Integrative Genomics Viewer [49]. Coverage over *env* sufficient for cladistics (Figure 4).

**Figure 2N:**
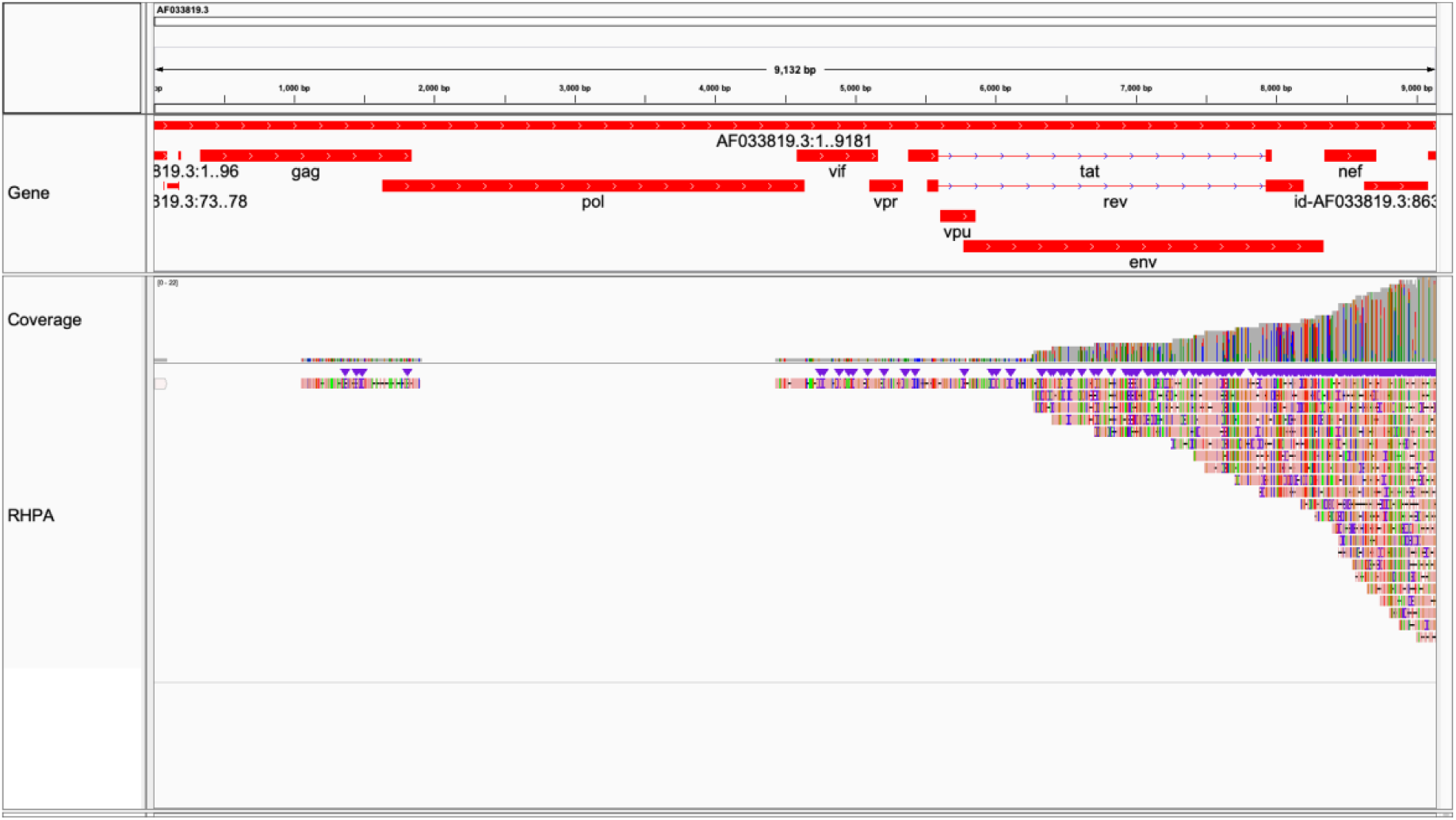
RHPA. Majority coverage (4,745/9,181∼51.7%) achieved. Visualized in Integrative Genomics Viewer [49]. Coverage over *env* insufficient for cladistics.

**Figure 2O:**
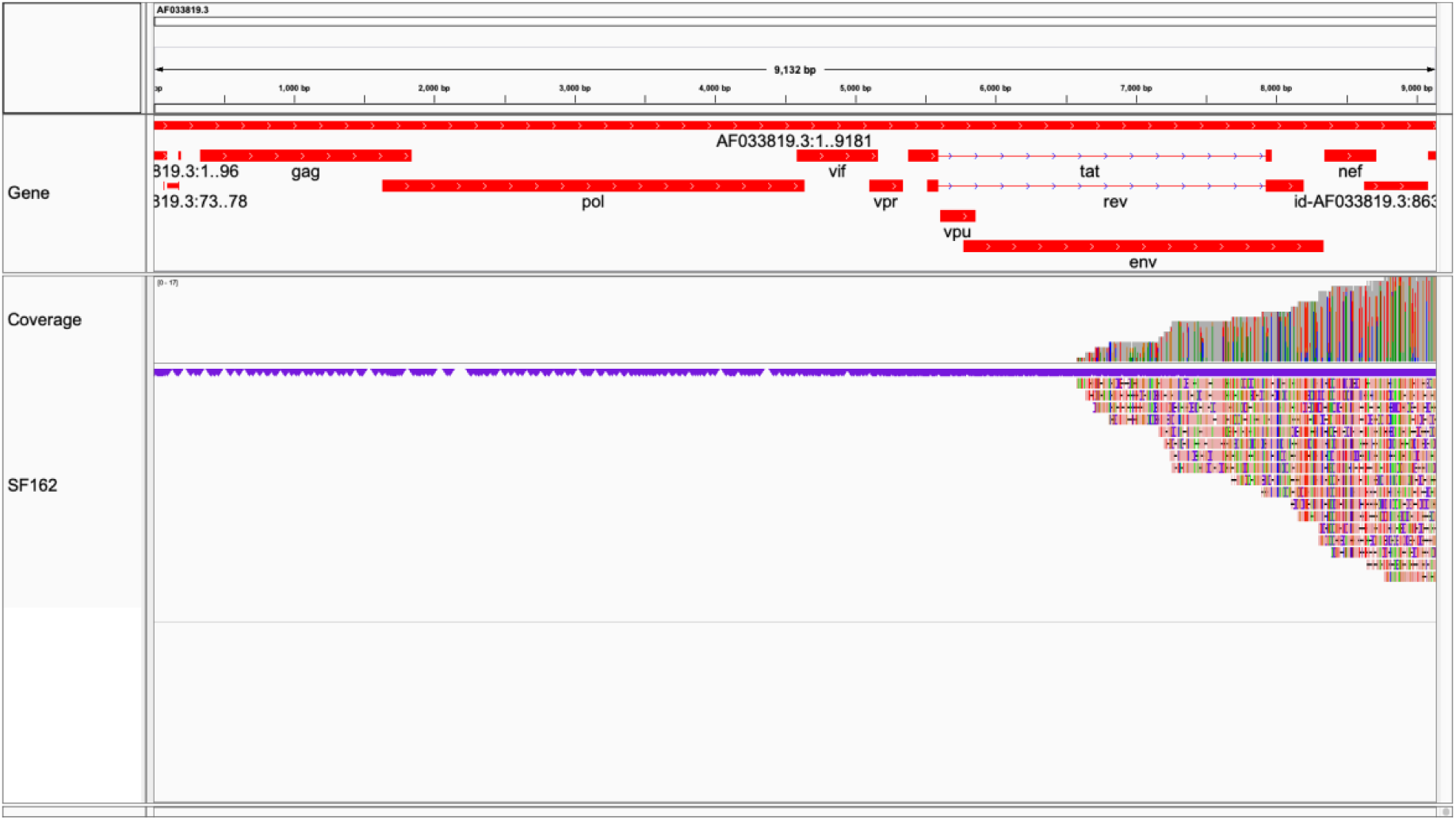
SF162. Moderate coverage (2,601/9,181∼28.3%) achieved. Note slight change in slope over RRE. Visualized in Integrative Genomics Viewer [49]. Coverage over *env* insufficient for cladistics.

**Table 1:**
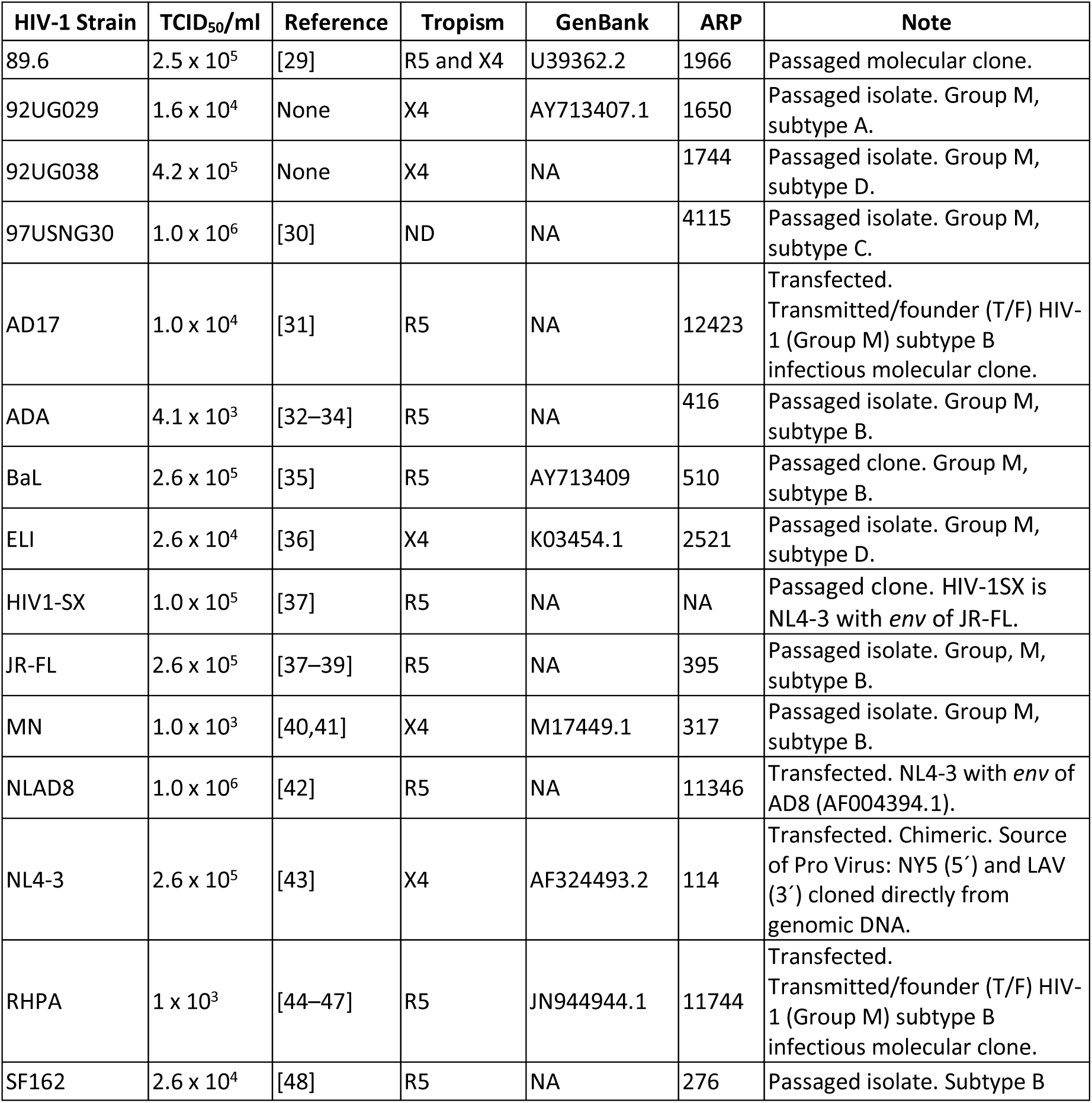
Fifteen HIV-1 strains used in this study. Abbreviations: TCID = Tissue culture Infectious Dose. R5 = CCR5. X4 = CXCR4. NA = not applicable. ND = not done. Contents of “Note” and “Tropism” columns summarized from the NIH AIDS Reagent and Reference Program (ARP; https://aidsreagent.org). GenBank accessions reported only for strains with complete genomes available.

| HIV-1 Strain | TCID<sub>50</sub>/ml | Reference | Tropism | GenBank | ARP | Note |
| --- | --- | --- | --- | --- | --- | --- |
| 89.6 | 2.5 x 10<sup>5</sup> | [29] | R5 and X4 | U39362.2 | 1966 | Passaged molecular clone. |
| 92UG029 | 1.6 x 10<sup>4</sup> | None | X4 | AY713407.1 | 1650 | Passaged isolate. Group M, subtype A. |
| 92UG038 | 4.2 x 10<sup>5</sup> | None | X4 | NA | 1744 | Passaged isolate. Group M, subtype D. |
| 97USNG30 | 1.0 x 10<sup>6</sup> | [30] | ND | NA | 4115 | Passaged isolate. Group M, subtype C. |
| AD17 | 1.0 x 10<sup>4</sup> | [31] | R5 | NA | 12423 | Transfected. Transmitted/founder (T/F) HIV-1 (Group M) subtype B infectious molecular clone. |
| ADA | 4.1 x 10<sup>3</sup> | [32–34] | R5 | NA | 416 | Passaged isolate. Group M, subtype B. |
| BaL | 2.6 x 10<sup>5</sup> | [35] | R5 | AY713409 | 510 | Passaged clone. Group M, subtype B. |
| ELI | 2.6 x 10<sup>4</sup> | [36] | X4 | K03454.1 | 2521 | Passaged isolate. Group M, subtype D. |
| HIV1-SX | 1.0 x 10<sup>5</sup> | [37] | R5 | NA | NA | Passaged clone. HIV-1SX is NL4-3 with *env* of JR-FL. |
| JR-FL | 2.6 x 10<sup>5</sup> | [37–39] | R5 | NA | 395 | Passaged isolate. Group, M, subtype B. |
| MN | 1.0 x 10<sup>3</sup> | [40,41] | X4 | M17449.1 | 317 | Passaged isolate. Group M, subtype B. |
| NLAD8 | 1.0 x 10<sup>6</sup> | [42] | R5 | NA | 11346 | Transfected. NL4-3 with *env* of AD8 (AF004394.1). |
| NL4-3 | 2.6 x 10<sup>5</sup> | [43] | X4 | AF324493.2 | 114 | Transfected. Chimeric. Source of Pro Virus: NY5 (5') and LAV (3') cloned directly from genomic DNA. |
| RHPA | 1 x 10<sup>3</sup> | [44–47] | R5 | JN944944.1 | 11744 | Transfected. Transmitted/founder (T/F) HIV-1 (Group M) subtype B infectious molecular clone. |
| SF162 | 2.6 x 10<sup>4</sup> | [48] | R5 | NA | 276 | Passaged isolate. Subtype B |

## RESULTS

### Coverage

HIV-1-mapping reads were recovered in all (n=15) experiments (**Figures 1**, 2). Three out of 15 strains had full-coverage, defined as read length spanning from the TSS to the 3’ LTR and mapped to AF033819.3. These were: 89.6 (9,166/9,181∼99.8%), AD17 (9,164/9,181∼99.8%), and NLAD8 (9,166/9,181∼99.8%). Seven out of 15 strains had majority coverage, defined as > 50% relative to AF033819.3. These were: 97USNG30 (6,992/9,181∼76.2%), BaL (8,597/9,181∼93.6%), ELI (6,987/9,181∼76.1%), MN (5,995/9,181∼65.3%), NL4-3 (4,715/9,181∼51.4%), RHPA (4,745/9,181∼51.7%), and HIV1-SX (5,561/9,181∼60.6%). Three out of 15 strains had moderate coverage < 50% and > 25% relative to AF033819.3. These were: 92UG038 (3,678/9,181∼40.1%), JR-FL (2,814/9,181∼30.7%), and SF162 (2,601/9,181∼28.3%). Two out of 15 strains had lower coverage <25% relative to AF033819.3. These were: 92UG029 (2,007/9,181∼21.9%), ADA (1,981/9,181∼21.6%). In general, samples with higher TCID_50_/ml were more likely to produce more and longer reads. Passed and failed reads were collapsed into per-sample datasets for this study based on the observation that ∼10% of reads could be recovered from failed read folders. Analysis of host-mapping RNA was deferred for this paper.

### 3’ bias

All experiments demonstrated strong 3’ coverage bias (**Figures 2**, 3). This was also seen in control ENO2 mRNA across all experiments (**Supplemental Figure 1**). The signal from the control template across experiments exhibited consistent decay in recovered read length. Evaluating the reduction in sequence across samples at consistent points (base 1,200 and 200) yielded an average loss of 65.55% (SD = 11.08) of recovered transcripts per 1,000 bases. (**Supplemental Table 2**). Furthermore, when we removed the outlier NL4-3 sequencing run from the analysis, which was completed with previously used but still functional R9.4 RevC flow cells and RNA002, the average loss was 68.41% (SD = 0.6521).

**Figure 3A:**
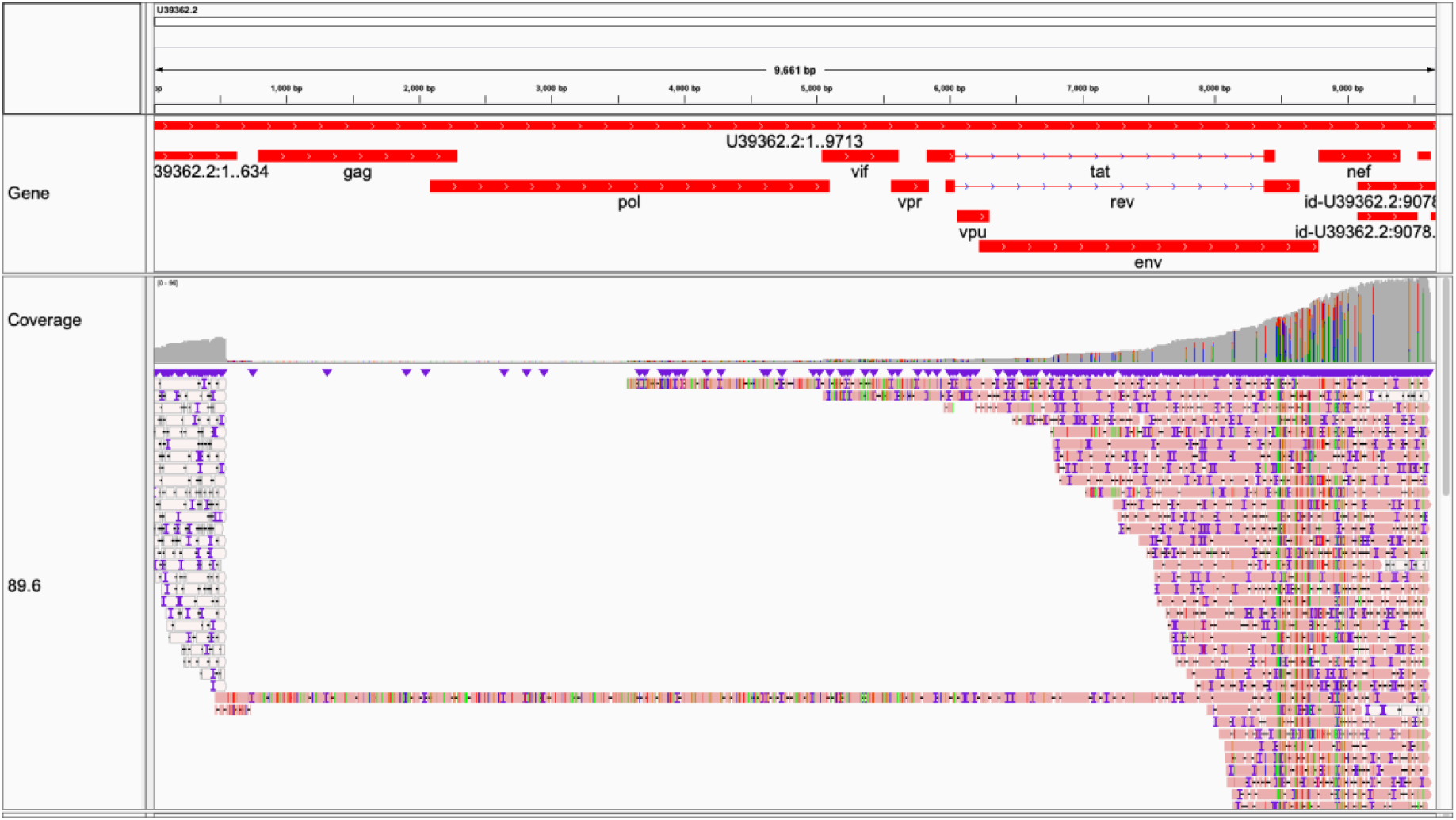
89.6 mapped to 89.6. Strain-specific reference mapping. Gray in coverage plot (and pink in read body) indicates per-base consensus accuracy ≥ 80%. Visualized in Integrative Genomics Viewer [49].

**Figure 3B:**
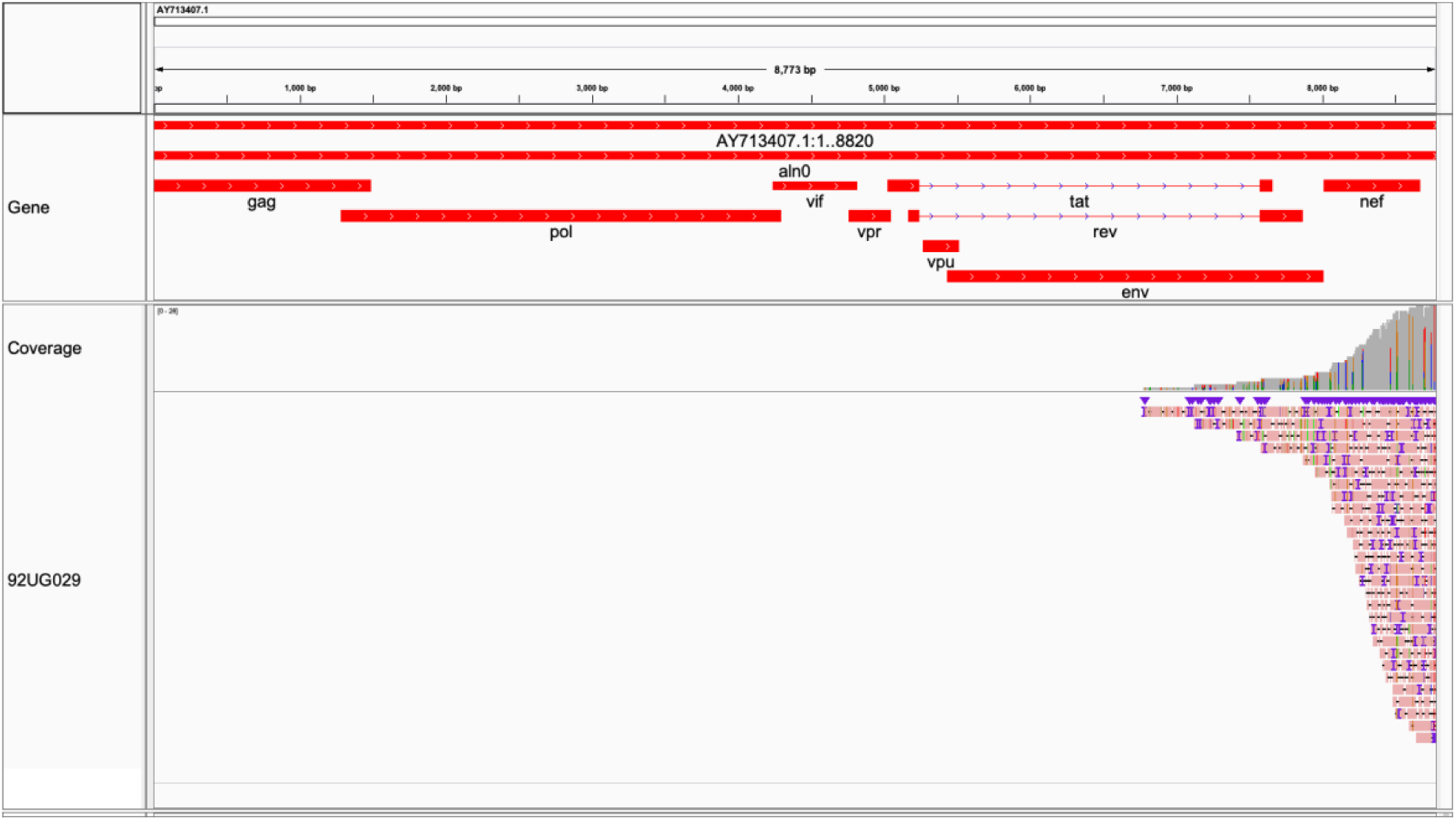
92UG029 mapped to 92UG029. Strain-specific reference mapping. Gray in coverage plot (and pink in read body) indicates per-base consensus accuracy ≥ 80%. Visualized in Integrative Genomics Viewer [49].

**Figure 3C:**
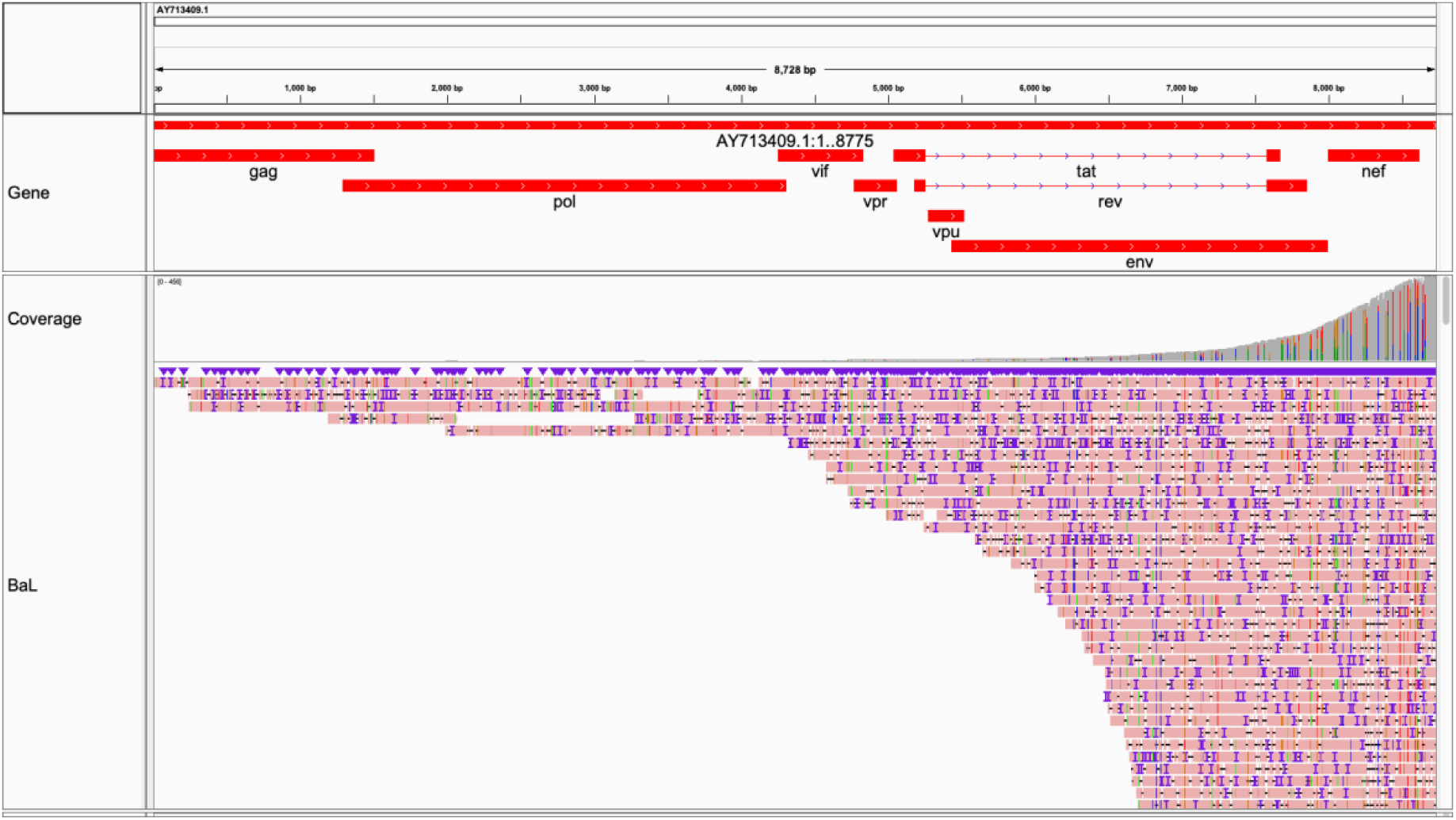
BaL mapped to BaL. Strain-specific reference mapping. Gray in coverage plot (and pink in read body) indicates per-base consensus accuracy ≥ 80%. Note BaL reference does not include LTRs. Visualized in Integrative Genomics Viewer [49].

**Figure 3D:**
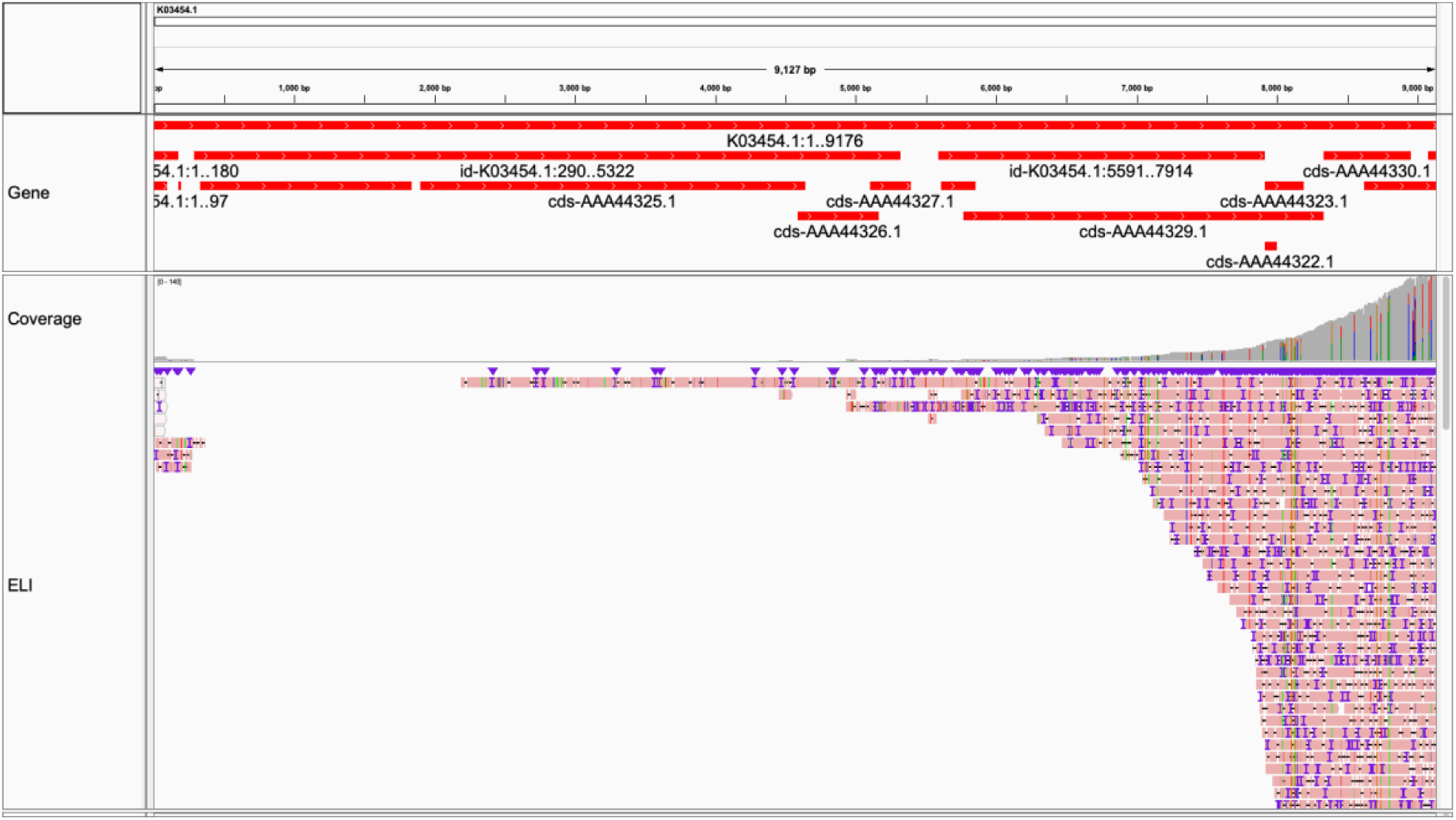
ELI mapped to ELI. Strain-specific reference mapping. Gray in coverage plot (and pink in read body) indicates per-base consensus accuracy ≥ 80%. Note slight change in slope over RRE. Visualized in Integrative Genomics Viewer [49].

**Figure 3E:**
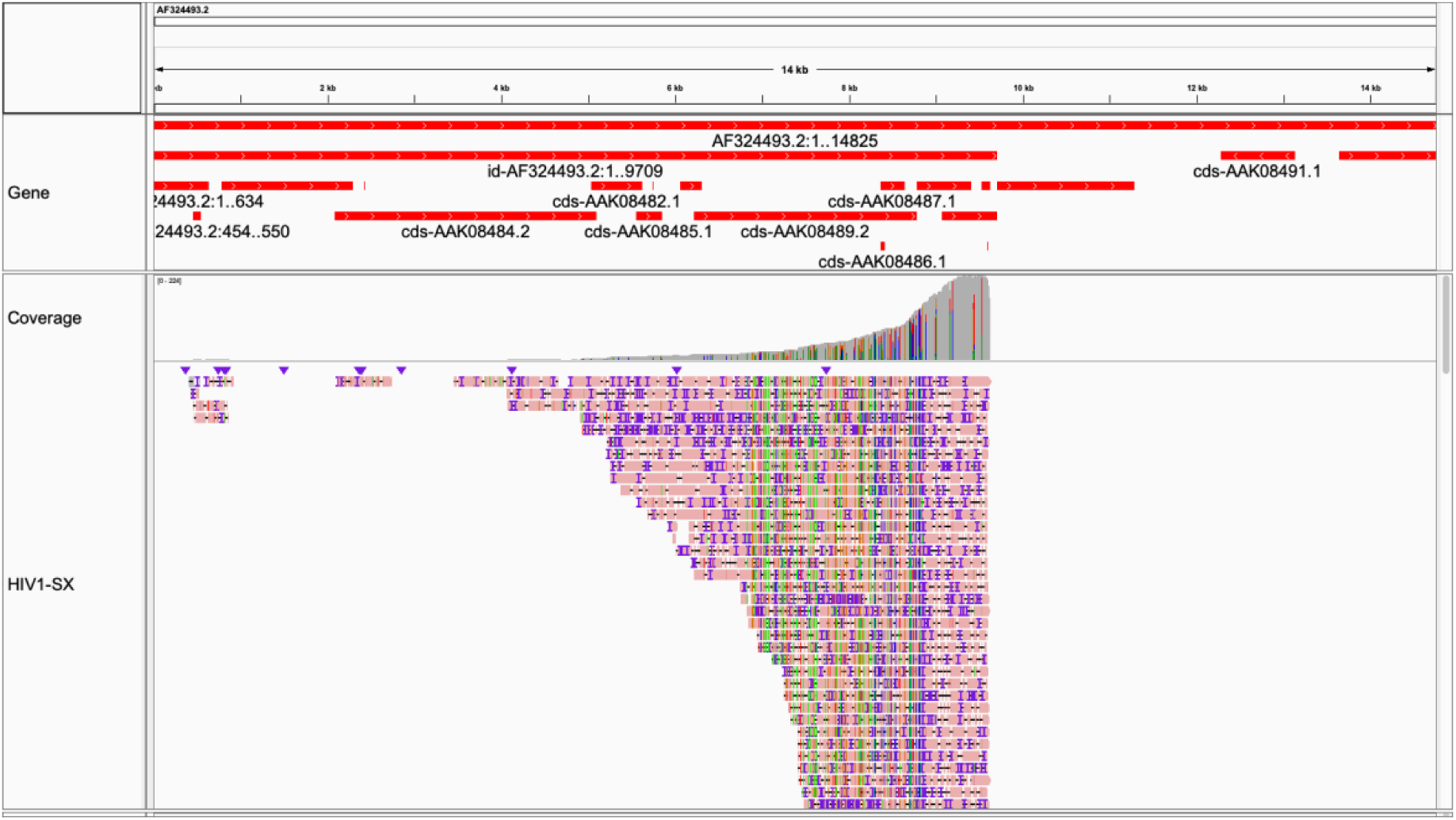
HIV1-SX mapped to pNL4-3. Strain-specific reference mapping. Gray in coverage plot (and pink in read body) indicates per-base consensus accuracy ≥ 80%. Visualized in Integrative Genomics Viewer [49].

**Figure 3F:**
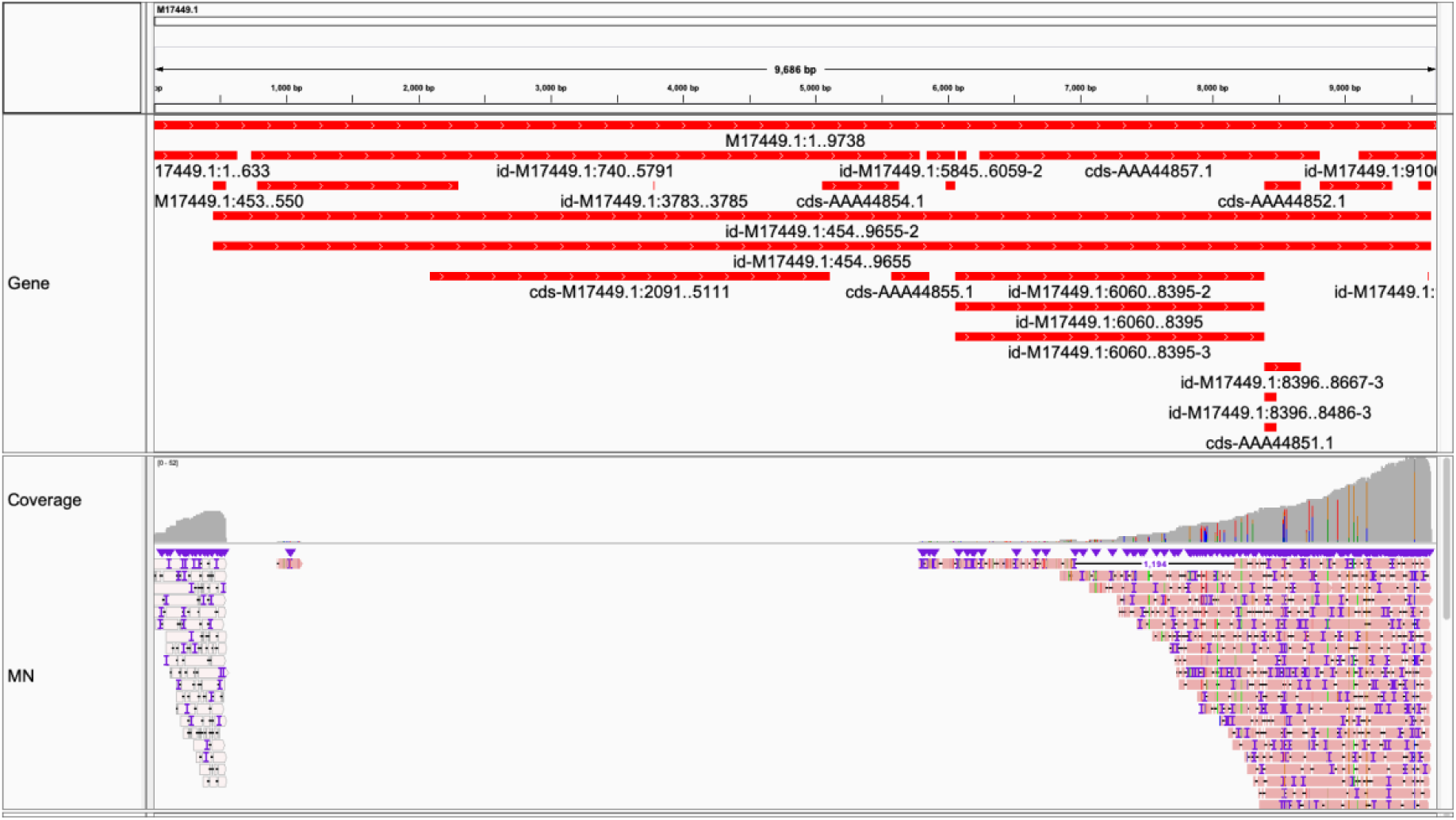
MN mapped to MN. Strain-specific reference mapping. Gray in coverage plot (and pink in read body) indicates per-base consensus accuracy ≥ 80%. Visualized in Integrative Genomics Viewer [49].

**Figure 3G:**
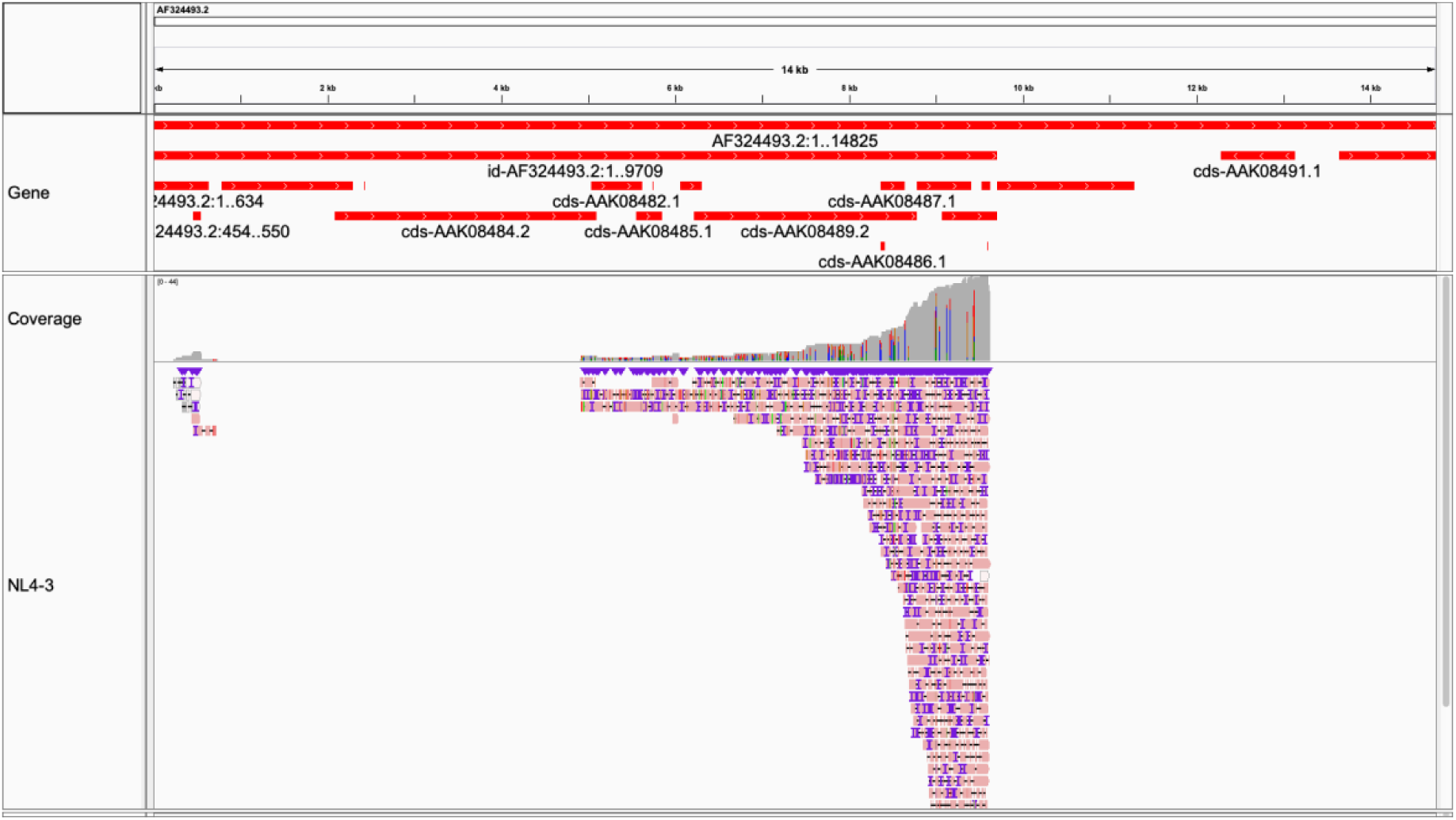
NL4-3 mapped to pNL4-3. Gray in coverage plot (and pink in read body) indicates per-base consensus accuracy ≥ 80%. Note slight change in slope over RRE. Visualized in Integrative Genomics Viewer [49].

**Figure 3H, 3I:**
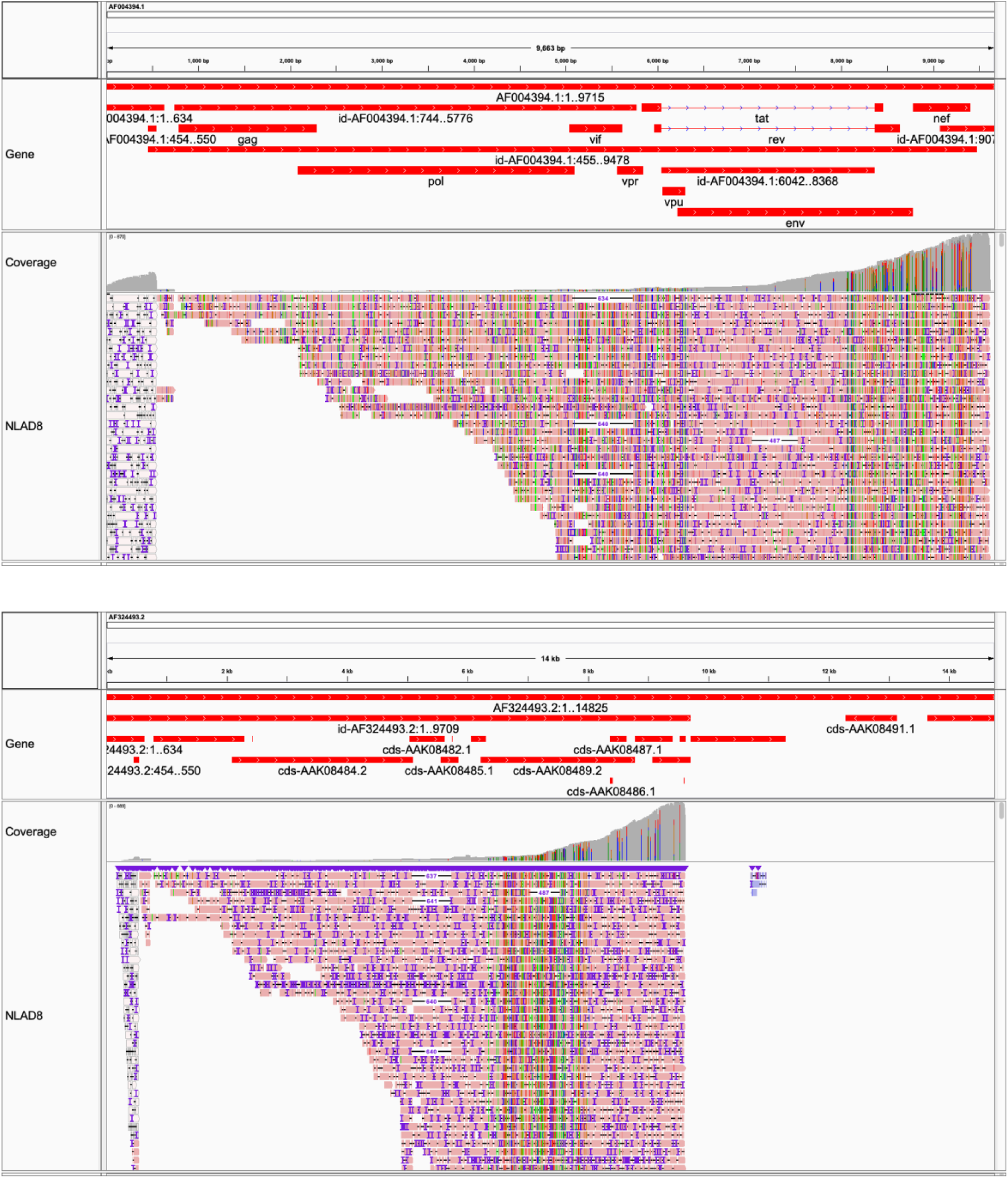
NLAD8 mapped to AD8, pNL4-3. Strain-specific reference mapping. Gray in coverage plot (and pink in read body) indicates per-base consensus accuracy ≥ 80%. Note that NLAD8 is a chimera between NL4-3 (all minus portion of *env*) and AD8 (portion of *env*) (used as a reference here). The reference sequence for NL4-3 is a plasmid, pNL4-3. NLAD8 complete genome was not available. Note the difference in scale. Visualized in Integrative Genomics Viewer [49].

**Figure 3J:**
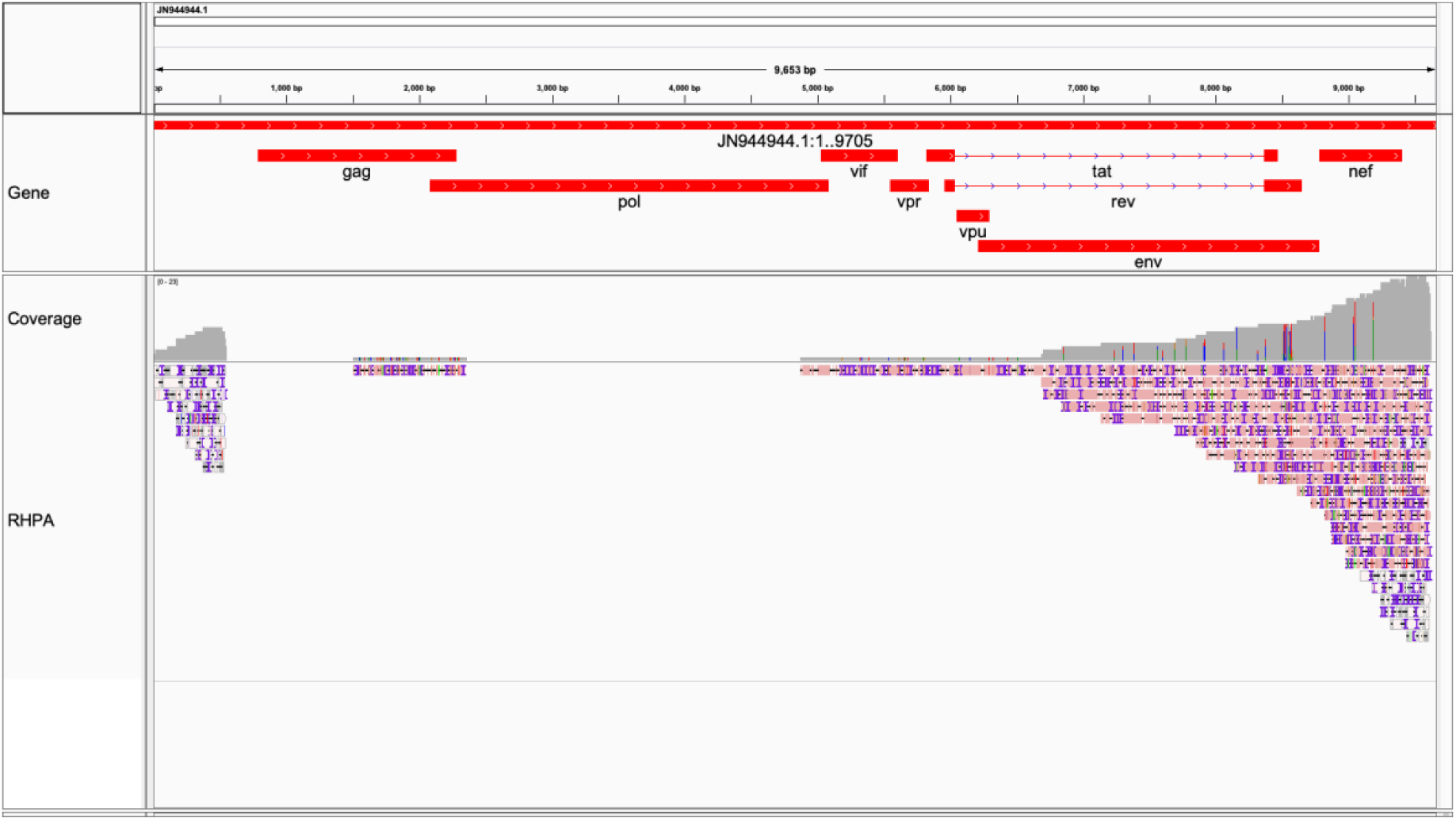
RHPA mapped to RHPA. Strain-specific reference mapping. Gray in coverage plot (and pink in read body) indicates per-base consensus accuracy ≥ 80%. Visualized in Integrative Genomics Viewer [49].

### Splicing vs. ONT artifact vs. Structural variant

We observed reads with large deletions in a higher coverage experiment (NLAD8 in **Figures 2E, 3H**, and **3I**). Without DNA sequencing, it is difficult to tell whether these are at the genomic or transcriptomic level. That 1) these were a minority, and that 2) the exact start and stop points differed between these few reads are suggestive of the latter. A smaller deletion was observed in a region of NLAD8 outside of *env* (**Supplemental Figure 5**). This small deletion had a different mapping profile compared to homopolymer runs (**Supplemental Figure 2**). The large deletions occurred at similar locations across multiple reads. Splicing was also evident as clipped reads in most samples (**Figure 2**). Other experiments either did not have such transcriptional features or lacked coverage sufficient to observe these features.

### SNV calls from ONT Native RNA

Samples with available reference genomes (with GenBank accessions) were manually evaluated for single nucleotide variants, including small insertions and deletions, under the conditions: 1) that there were at least 10 reads at a given locus; and 2) that the reference allele occurred less than or equal to 50% of the time. In the event that a putative SNV occurred over a deletion, the SNV was included in the set. **Supplemental Figures 4-7** include SNVs, small insertions, deletions. See **Supplemental Table 3** for a list of likely SNVs directly called from native (ONT) RNA-seq.

The region satisfying the above for 89.6 was 6,831-9,624 (length = 2,793) with respect to the reference 89.6 U39362.2. There were at least 45 SNVs in the version of 89.6 sequenced. The mutation rate, calculated as the number of SNVs divided by window length, was 45/2,793 = 0.01611171. The region satisfying the above for 92UG029 was 8,154-8,820 (length = 666) with respect to its reference AY713407.1. There were at least 7 SNVs in the version of 92UG029 sequenced. The mutation rate was 7/666 = 0.01051051. The region satisfying the above for BaL was 4,733-8,775 (length = 4,042) with respect to its reference AY713409.1. There were at least 23 SNVs in the version of BaL sequenced. The mutation rate was 23/4,042 = 0.00569025. The region satisfying the above for ELI was 7,105-9,176 (length = 2,071) with respect to its reference K03454.1. There were at least 18 SNVs in the version of ELI sequenced. The mutation rate was 18/2,071 = 0.00869145. The region satisfying the above for HIV1-SX was 5,385-9,624 (length = 4,239) with respect to a related reference pNL4-3 AF324493.2. However, HIV1-SX is chimeric, with JR-FL’s *env* region. Excluding *env*, there were no discernable differences between HIV1-SX and pNL4-3. The region satisfying the above for MN was 7,752-9,654 (length = 1,902) with respect to its reference M17449.1. There were at least 11 SNVs in the version of MN sequenced. The mutation rate was 11/1,902 = 0.00578339. The region satisfying the above for NL4-3 was 8,169-9,625 (length = 1,456) with respect to its reference AF324493.2. There were at least 2 SNVs in the version of NL4-3 sequenced. The mutation rate was 2/1,456 = 0.00137363. The region satisfying the above for NLAD8 was 1,474-9,626 (length = 8,152) with respect to its reference AF324493.2. There were at least 6 SNVs in the version of NLAD8 sequenced. NLAD8 was completely conserved (no obvious SNVs) in its *env* region compared to AD8 (**Figure 3H, Figure 3I**). The region satisfying the above for RHPA was 8,324-9,621 (length = 1,297) with respect to its reference JN944944.1. There was one SNV in the version of RHPA sequenced, g.8,570C>T, nonsynonymous T>M in *rev* and nonsynonymous R>C in *env*, with possible structural consequence. The mutation rate was 6/8,152 = 0.00073602. Combined, the average mutation rate (excluding HIV-SX and RHPA) was 0.00698528, (SD = 0.00535872). We noticed variants occurring in close proximity to other neighboring variants. These represent interesting cases of context-dependent variance, which we classify with the following system: Independent, Neighbor, Neighborhood, or Neighbor+Neighborhood to emphasize the points that adjacent bases can contribute to multiple amino acid changes, and that these are not immediately apparent when focusing on individual coding frames (See **Supplemental Figure 8**).

### Comparative transcriptomics

HIV-1-mapped (AF033819.3 with minimap2) reads were assembled into contigs with Canu, a leading *de novo* long read genome assembler [10]. Contigs were generated for 89.6, 97USNG30, AD17, BaL, ELI, HIV1-SX, MN, NLAD8 (**Supplemental Figures 9, 10**). Contigs were fed into MAFFT server [15], and basic phylogenetic trees generated with default settings (**Figure 4**). For sequences with GenBank accessions with complete genomes, the tree was homogenous. With incomplete contigs, general relationships (such as similarity to NL4-3) were maintained. By mapping reads to references we observed that phasing (the ability to tie SNVs to individual viral chromosomal haplotypes) was possible with native RNA reads (**Supplemental Figure 7**). We wanted to test whether this could be leveraged to study viral transcript heterogeneity directly. All FASTQ were converted to FASTA, concatenated, made into a multisequence alignment, and visualized as a tree (**Figure 4**). Surprisingly, sample relationships were maintained despite per-read noise.

**Figure 4:**
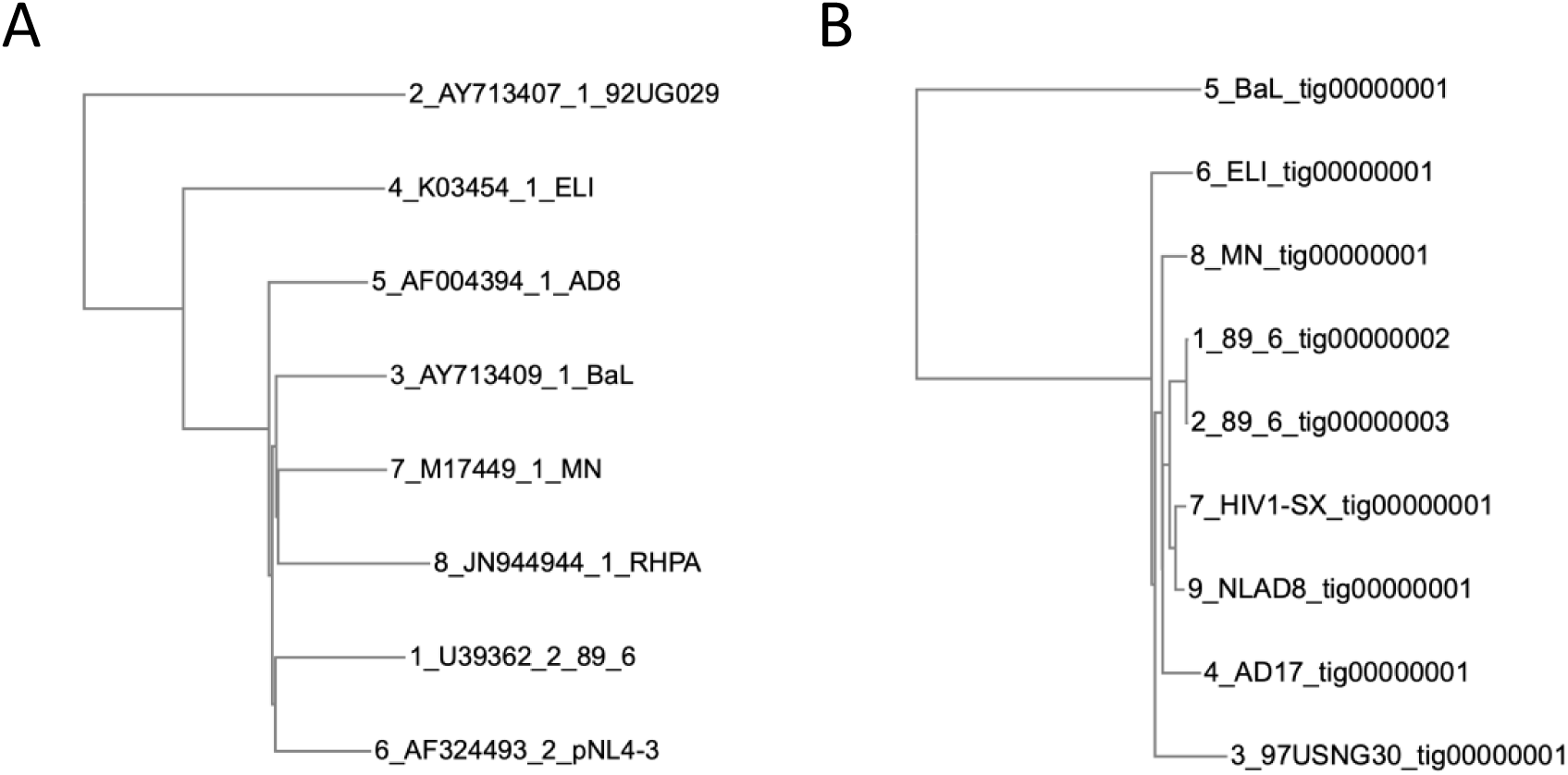
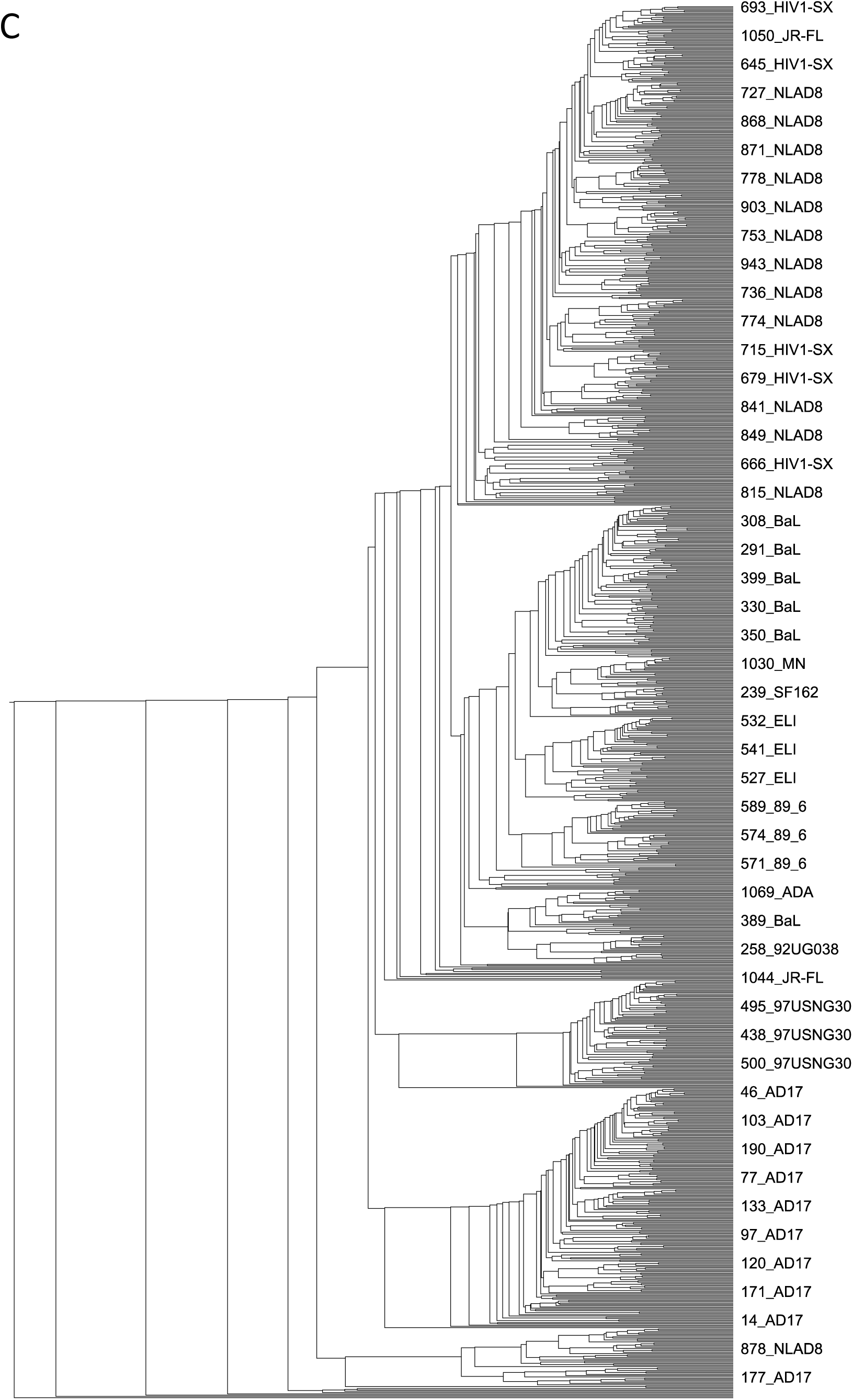
ONT native RNA + Canu *de novo* assembly sufficient for comparative transcriptomics. Trees from multisequence alignments. Phylogenetic relationships (subtype, relationship to NL4-3) evident in trees from full reference genomes are generally conserved at the contig and individual sequence levels. A. Available full-length genomes (left). B. Contigs (right). C. All recovered HIV reads. MAFFT [15].

## DISCUSSION

### First complete full-length sequences from infectious HIV-1 strains

In this study we were able to completely [16] sequence the viral RNA genome of three HIV-1 strains (**Figure 1**). Here, we defined complete or full-coverage of full-length HIV-1 viral RNA as the region capable of sustaining viral replication with supplemental reverse transcription and integrase activity. This corresponds to the region from the 5’ LTR-mediated transcriptional start site (TSS) and the repeat region after the end of the U3 region of the 3’ LTR. Note that these dimensions are different than the dimensions of HIV-1 provirus. To our knowledge, this is the first time the HIV-1 virus has been sequenced from the TSS to end of 3’ LTR. As reported for HIV-1 in a DNA context [3], ONT surpassed the read length limitations of short read RNA-seq. Data were sufficient for SNV detection over regions with coverage ≥ 10 reads, as well as direct *de novo* assembly from native RNA reads, with a maximal assembly length of 94% from strain NLAD8.

### Mapping

Historically lower per-read accuracy and high sequence variability across HIV isolates between and within patients represent two challenges to detecting native HIV reads. Interestingly, newer long read mappers such as minimap2 were robust to references used (Mapping quality >50). We saw slightly more recovered reads when we mapped our reads to references with greater homology (**Figures 2A** and **3A; Figures 2C** and **3B; Figures 2G** and **3C; Figures 2H** and **3D; Figures 2K** and **3F; Figures 2L** and **3G; Figures 2M, 3H and 3I; Figures 2N** and **3J**). We recommend using strain-specific references when possible, ideally sequence-verifying constructs used to make viruses in each lab to catch any lurking SNVs [3].

### Coverage

These experiments represent a proof-of-concept demonstrating ONT’s ability to cover HIV mRNA completely (**Figures 2A, 2E, 2M, 3A, 3H**, and **3I**), dependent on the titer of viruses used for library prep (**Table 1**). Neither cDNA nor cDNA PCR ONT libraries were evaluated in the present study. Coverage as read depth (total number of bases sequenced divided by reference length) was not informative due to strong 3’ bias.

### 3’ bias

We were able to recover full-length HIV RNA reads from 3 samples. This was in part due to strong 3’ bias inherent to the RNA002 library. The first adapter used by the RNA002 kit, RTA, selects for polyA [7]. There was relatively consistent coverage exhibiting a decrescendo pattern (with coverage decreasing distal from RTA ligation point), except near areas of known complex 2° RNA structure [17]. The first strand cDNA step is supposed to ameliorate 2° RNA, but these might not be completely abrogated—for example, the region around the rev response element in **Figure 2, 3** in higher-coverage experiments exhibited slight changes in average loss of recovered transcripts per 1,000 bases. A useful consequence of the 3’ bias is easier phasing between 5’ and 3’ LTR regions, which is difficult or impossible to tell apart using short read DNA [3] or classic RNA sequencing (**Supplemental Figure 12**). In fact, we did not encounter reads spanning full-length 5’ LTR in any of our native RNA experiments, which were however present in HIV PCR + DNA ONT experiments [3].

Included in the RNA002 library kit is control RNA. By determining the percent loss per 1,000 bases, we recommend that at least 2,500x coverage over the 3’LTR to guarantee observation of full-length HIV reads (∼9 kb) at a depth sufficient to call SNVs at the 5’ end of transcripts (with coverage ∼60x, which we recommend for Canu). At current sequencing throughput, this is equivalent to about one MinION RevD flow cell, with possibly multiple technical replicates of individual libraries. Note that higher-titer virus stocks were associated with higher coverage, although the sequencing time was not held constant.

### Splicing vs. ONT artifact vs. Structural variant

In NLAD8 (**Figures 2M, 3H**, and **3HI**), there was high enough coverage toward the middle of HIV to see deletions relative to reference. These read deletions might be splicing, artifacts from library prep (with T4 DNA ligase), or due to previously observed HIV-1 deletion-prone integrants [18–20]. Note that these deletions are individual observations and not necessarily representative of the strains sequenced. A possible explanation supporting splicing includes observations of splicing events in a short read (cDNA) RNA-seq experiment (collected by Siarhei Kharytonchyk and submitted by Matthew Eckwahl). This incompletely published experiment was most likely using virus derived from pNL4-3, a commonly used lab-adapted clone of HIV-1 (Gener, unpublished). (See **Supplemental Figure 12**.) Mapping peaks also occurred at sites of known exons. On closer inspection, these reads exhibited clipping indicating their being part of a spliced read.

### SNV calls from ONT Native RNA

Strain-specific reference mapping was performed on 89.6, 92UG029, BaL, ELI, MN, NL4-3, and NLAD8. Abundance of coverage allowed us to evaluate single nucleotide variants (SNVs), as well as small insertions and deletions, in areas with ≥ 10x coverage (minimum recommended by Canu documentation). The average mutation rate across sequenced samples (**Supplemental Table 3**) was 0.00698528. This meant that for each full-length HIV-1 virion (say with respect to the HIV-1 reference AF033819.3) one would expect 0.00698528 × 9,181 bases = ∼64 SNVs, possibly contributing to altered protein products. We introduce neighbor and neighborhood nomenclature for SNVs to emphasize that variants occurring in information-dense regions can have complex context-dependent effects. What was demonstrated for HIV in its DNA form [3] also applies for HIV in its RNA form: SNVs happen. Sequencing the viral strains and cloned constructs used for experiments in a given laboratory remains important, because what is reported in a reference database for a given virus may not necessarily be exactly the same as the viruses propagated in a laboratory.

### Capturing variability in reference HIV-1 viruses

HIV is the most studied human pathogen [3], and yet there were at least 144 SNVs across 9 HIV-1 reference strains. Some occurred in previously camouflaged [21] repetitive LTR regions, but most occurred in the gene bodies of sequenced strains, some contributing to alterations in protein-coding potential. In this work we leveraged sequencing multiple related strains with mapping to their predecessors to simulate true positive reads. Long read native RNA sequencing on the ONT MinION with R9.4.1 RevD flow cells was robust and sufficient to detect real sequence variability at the single-molecule level, except for at areas in and around homopolymers. While not a complete solution yet, long read native RNA sequencing offers the ability to tie SNVs together, to define segments of HIV haplotypes directly, and to move away from the concept of quasispecies [1, 22] toward defining real viral haplotypes. An extension of this capability is the ability to perform comparative transcriptomics within hours after sample acquisition (**Figure 4**), including determining drug resistance profiles based on existing public databases (**Supplemental Figure 11**) [23, 24].

### Homopolymers

As previously reported [3, 25] for DNA ONT, per-read variability in ONT data was higher near homopolymers (runs of the same base) (**Supplemental Figure 2)**. ONT is also known for lower per-base accuracy compared to short read next generation platforms. These may be overcome by evaluating neighbor SNVs (**Supplemental Figure 3**). New developments include improved basecalling models and a new double-header R10 nanopore, which has been showing improved handling of homopolymers, with slight drop in per-read accuracy. When comparing sequences, the errors in homopolymers are observed consistently across reads, and become negligible. We were able to compare individual reads from mostly incomplete virion genomes with simple cladistics and reads successfully grouped based on sample identity (**Figure 4**).

### Suitability of ONT for HIV sequencing

Comparing Classic RNA-seq to Native RNA-seq, each technology has its pros and cons. Classic RNA-seq, usually cDNA synthesis optionally followed by PCR, can be sensitive, specific, and cheaper with newer higher throughput sequencers (example: Illumina, BGI). While PCR enhances sensitivity (to this day, it is important for detecting viral sequences at low copy numbers [26, 27]), a downside is that captured information can be lost if PCR is used during library prep (**Supplemental Figure 12**). Direct cDNA, or using long read DNA sequencing (PacBio [2], ONT), can also recuperate sequence information from samples with relatively higher throughput than native RNA-seq in its current implementation. It will be important to address sequencing throughput and sequence biases when developing this nascent technology. At present, it is possible to sequence and compare viral isolates as contigs and for the first time as individual virus genomes as individual reads (**Figure 4**), as well as to determine drug resistance profiles from contigs (**Supplemental Figure 11**) or individual reads. A benefit that is unique to native RNA sequencing on ONT devices is the ability to detect base modifications as well. Work is ongoing to evaluate the raw signal from these experiments, and to evaluate new ONT RNA analysis tools for use with HIV.

### Closing the distance on HIV-1 with longer reads

Current limitations of the approaches used in the present work to study HIV RNA are similar to those reported for HIV DNA [3], and include: 1) the cost of long-read sequencing, compared to the cheaper short read DNA sequencing (as in classic RNA-seq); 2) long RNA extraction methods in diseased tissue (Gener, unpublished); and 3) lower per-base accuracy (mid 90’s vs. 98-99%), including difficulty near homopolymers (**Supplemental Figure 2**). Furthermore, a unique major limitation to RNA sequencing is 4): 3’ bias. As the price of long-read sequencing continues to decrease, and as the technology improves, the cost of obtaining usable data from native RNA long read sequencing will become negligible compared to the ability to answer new questions. Classic RNA-seq had a problem with 3’ bias, but the issue was eventually overcome [28]. Future work will move toward the goal of capturing higher-coverage fuller glimpses of each HIV viral mRNA, including virion genomes in *in vivo* HIV models and from patient samples. Long read sequencing is an important emerging tool defining the post-scaffold transcriptomic era, allowing for the characterization of functional units at the intersection between host and pathogen transcriptomes.

## ACKNOWLEDGEMENTS

ARG conceived of this project and performed all experiments. ARG and JK analyzed results and wrote the manuscript. The authors would like to thank Dr. Sue Ellen Crawford for earlier involvement and Victoria Rose Tenge for advice, technical assistance, and critical reading of the manuscript.

This work was funded in part by institutional support from Baylor College of Medicine; the Human Genome Sequencing Center, BCM; private funding by East Coast Oils, Inc., Jacksonville, Florida, and ARG’s own private funding, including Student Genomics (manuscripts in prep). Compute resources from the Computational and Integrative Biomedical Research Center at BCM (“sphere” cluster managed by Dr. Steven Ludtke) and the Department of Molecular and Human Genetics at BCM (“taco” cluster managed by Mr. Tanner Beck, Mr. Christopher Michael Holder, and Dr. Charles Lin) greatly facilitated the completion of this work. ARG has also received the PFLAG of Jacksonville scholarship for multiple years. The authors would like to thank Dr. David Raul Murdock in Dr. Brendan Lee’s Laboratory and Mr. Alexander Robert Kneubehl in Dr. Job E. Lopez’s Laboratory for making their MinION devices available. The authors would like to thank the Mary K. Estes Laboratory at BCM for making available their Kingfisher instrument for viral RNA isolation. This work was also supported in part by NIH grant AI116167 to JTK.

Infectious virus stocks of HIV-1 (SF162, 97USNG30, MN, ADA, 92UG038, 92UG029, ELI, JR-FL, and BaL) or molecular clones (NL4-3, NLAD8, RHPA, AD17) were obtained from M. Martin, R. Collman, E.O. Freed, B. Hahn, G. Shaw, J. Levy, D. Ellenberger,,P. Sullivan, R.B. Lal, R.C. Gallo, H. Gendleman, I.S.Y. Chen, S. Gartner, M. Popovic, and the UN AIDS Network for HIV Isolation and Characterization and DAIDS, NIAID via the NIH AIDS Reagent and Reference Program. HIV1-SX was provided by M.R. Ferguson and W.A. O’Brien.

ARG would like to thank members of the Paul E. Klotman Laboratory at Baylor College of Medicine, including Dr. Paul E. Klotman, Dr. Deborah P. Hyink (who thoughtfully helped to edit the manuscript), Taneasha Washington, and former members Dr. Gokul C. Das and Alexander Batista.

This work is dedicated to the memory of Dr. James “Jim” E. Maruniak, an outstanding virologist, educator, family man, role-model, and friend.

## SUPPLEMENTAL DATA

**Supplemental Table 1:** Library Statistics.

| Sample | Read count | Average length | Median length | Max length | Avg mean QS | Median mean QS |
| --- | --- | --- | --- | --- | --- | --- |
| 89.6 | 50 | 1258 | 922 | 8679 | 8.31 | 8.56 |
| 92UG029_run1 | 4 | 1161 | 1116 | 1731 | 10.42 | 10.45 |
| 92UG029_run2 | 2 | 937 | 937 | 989 | 9.08 | 9.08 |
| 92UG029_run3 | 2 | 1180 | 1180 | 2023 | 9.66 | 9.66 |
| 92UG038 | 4 | 1159 | 1240 | 1401 | 10.19 | 9.76 |
| 92UG038_skip | 8 | 1622 | 1490 | 3678 | 10.63 | 10.71 |
| 97USNG30 | 22 | 1333 | 1151 | 3742 | 10.56 | 10.48 |
| 97USNG30_skip | 61 | 1292 | 1007 | 4400 | 10.96 | 11.09 |
| AD17_run1 | 42 | 2026 | 1508 | 7176 | 10.08 | 10.12 |
| AD17_run1_skip | 109 | 1760 | 1343 | 5508 | 10.30 | 10.48 |
| AD17_run2 | 31 | 2303 | 1700 | 7390 | 9.79 | 10.03 |
| AD17_run2_skip | 12 | 2249 | 1256 | 8830 | 10.52 | 10.71 |
| ADA_run1 | 5 | 617 | 565 | 845 | 9.79 | 9.77 |
| ADA_run2 | 1 | 1334 | 1334 | 1334 | 11.14 | 11.14 |
| ADA_run3 | 4 | 852 | 616 | 1904 | 9.58 | 9.90 |
| ADA_run4 | 0 | NA | NA | NA | NA | NA |
| BaL | 22 | 969 | 905 | 2038 | 9.79 | 9.81 |
| BaL_skip | 147 | 1020 | 765 | 4474 | 10.08 | 10.26 |
| ELI_run1 | 9 | 2017 | 809 | 6909 | 10.08 | 10.11 |
| ELI_run1_skip | 30 | 905 | 763 | 2887 | 9.72 | 10.01 |
| ELI_run2 | 5 | 1017 | 989 | 1921 | 11.05 | 11.62 |
| ELI_run2_skip | 7 | 890 | 531 | 1892 | 9.98 | 10.03 |
| HIV1-SX | 107 | 1278 | 954 | 4796 | 10.02 | 10.10 |
| JR-FL | 21 | 1345 | 783 | 4695 | 9.98 | 9.75 |
| MN | 25 | 846 | 638 | 1912 | 10.12 | 10.41 |
| NL4-3_run3 | 19 | 1211 | 948 | 4626 | 9.92 | 10.23 |
| NL4-3_run5 | 9 | 1475 | 1021 | 4630 | 9.04 | 8.80 |
| NLAD8 | 291 | 1577 | 1081 | 8956 | 9.97 | 10.16 |
| RHPA | 13 | 1447 | 890 | 4696 | 10.39 | 10.90 |
| SF162 | 6 | 1849 | 1950 | 2540 | 10.60 | 11.34 |
| SF162_skip | 5 | 1041 | 862 | 2020 | 10.48 | 10.20 |

**Supplemental Figure 1:**
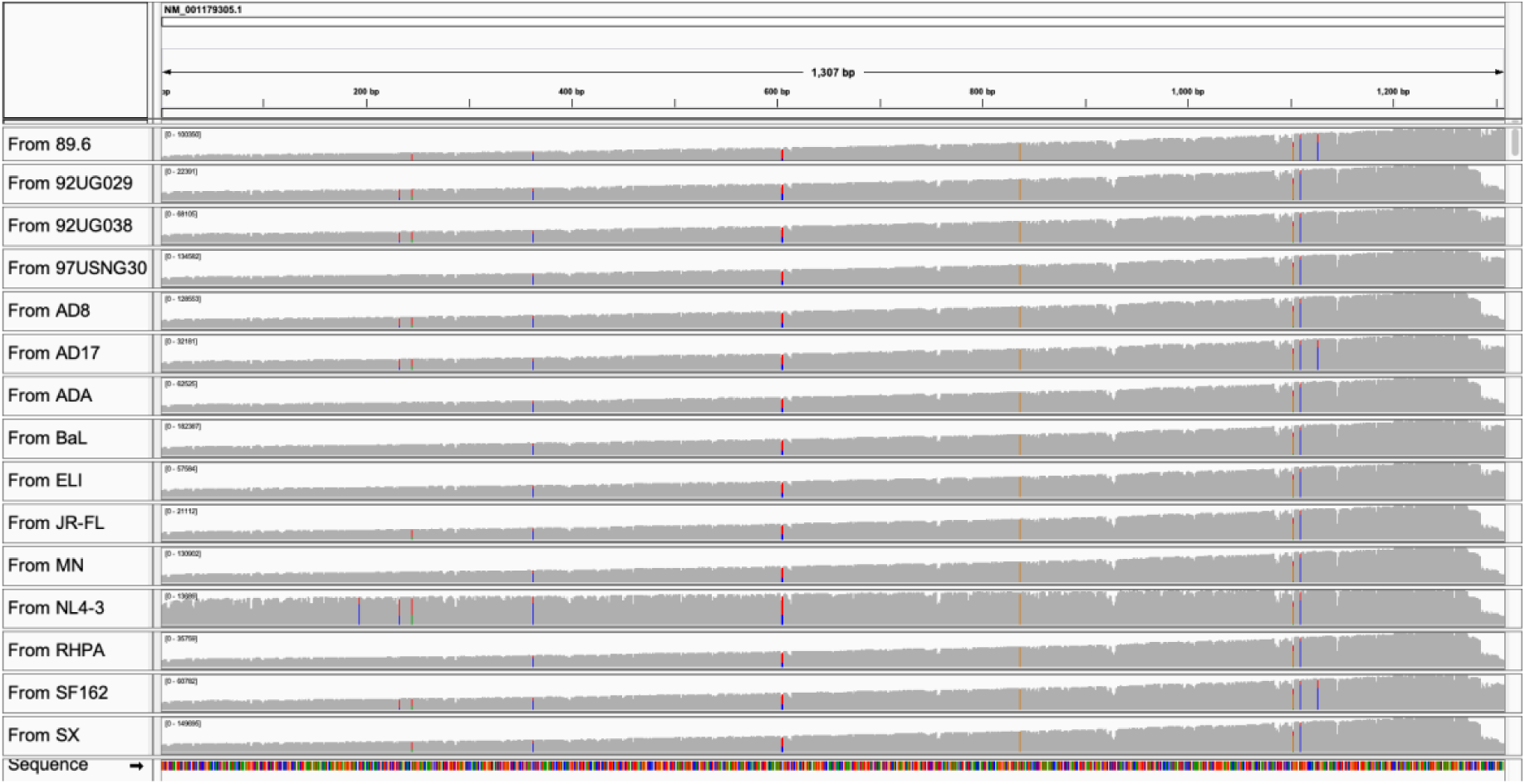
Consistent mapping and read quality between biological replicates. NL4-3 was processed with a previously used R9.4 RevC flow cell (now discontinued) in a prior pilot study. All other samples processed with R9.4.1 RevD flow cells. See **Methods** section for more details about control RNA. Alignments with minimap2 [13] in usegalaxy.eu [14]. Visualized in Integrative Genomics Viewer [49].

**Supplemental Table 2:**
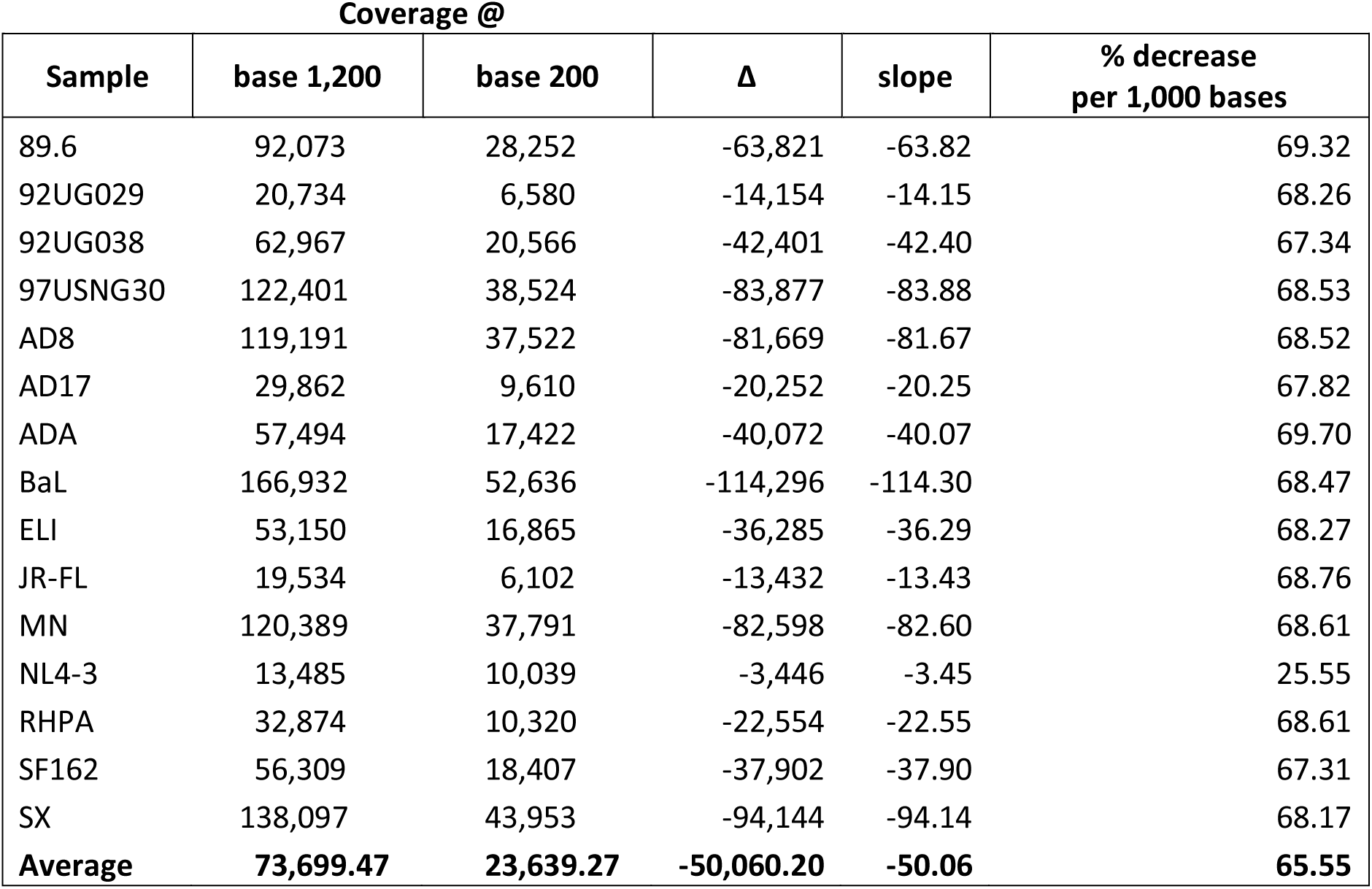
For SQK-RNA002, coverage drops by approximately 2/3rds per 1,000 bases. Data from Control RNA, which has much higher coverage compared to viral or host cell RNA. Positions relative to reference. Counts from reads mapped to ENO2 (minimap2 in Galaxy), visualized in Integrative Genomics Viewer [49].

**Supplemental Figure 2:**
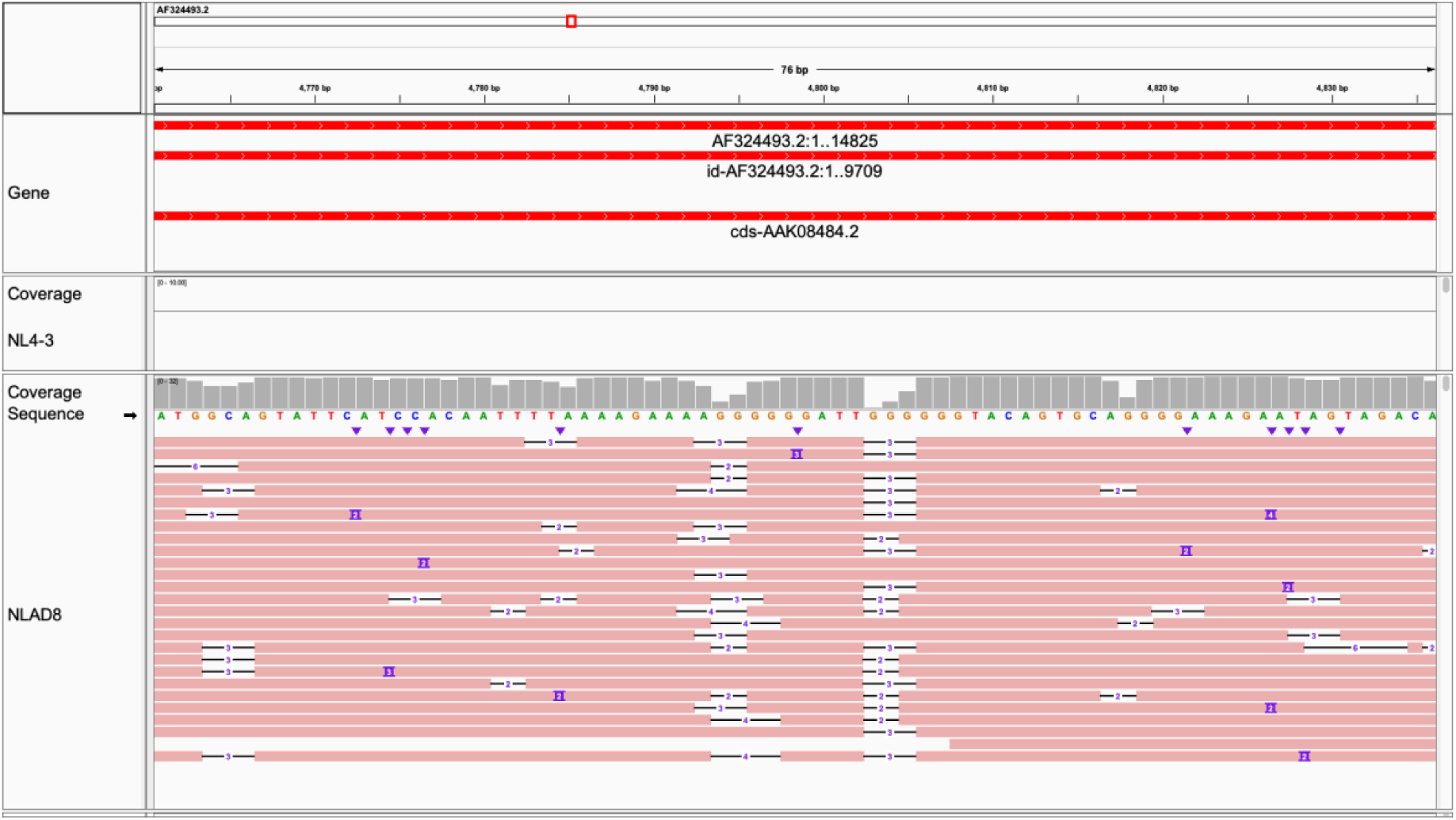
A set of homopolymer tracks from HIV-1 NLAD8. Top: AF324493.2:4,761-4,836. Alignments with minimap2 [13] in usegalaxy.eu [14]. Bottom: AF033819.3: 4,308-4,383. Note absence of insertions near trailing ends of homopolymers, supporting mapping as an important QC step when calling variants near these sequence features. Visualized in Integrative Genomics Viewer [49]. Comparable to DNA ONT [3]. The native NL4-3 sample did not have coverage over this region.

**Supplemental Figure 3:**
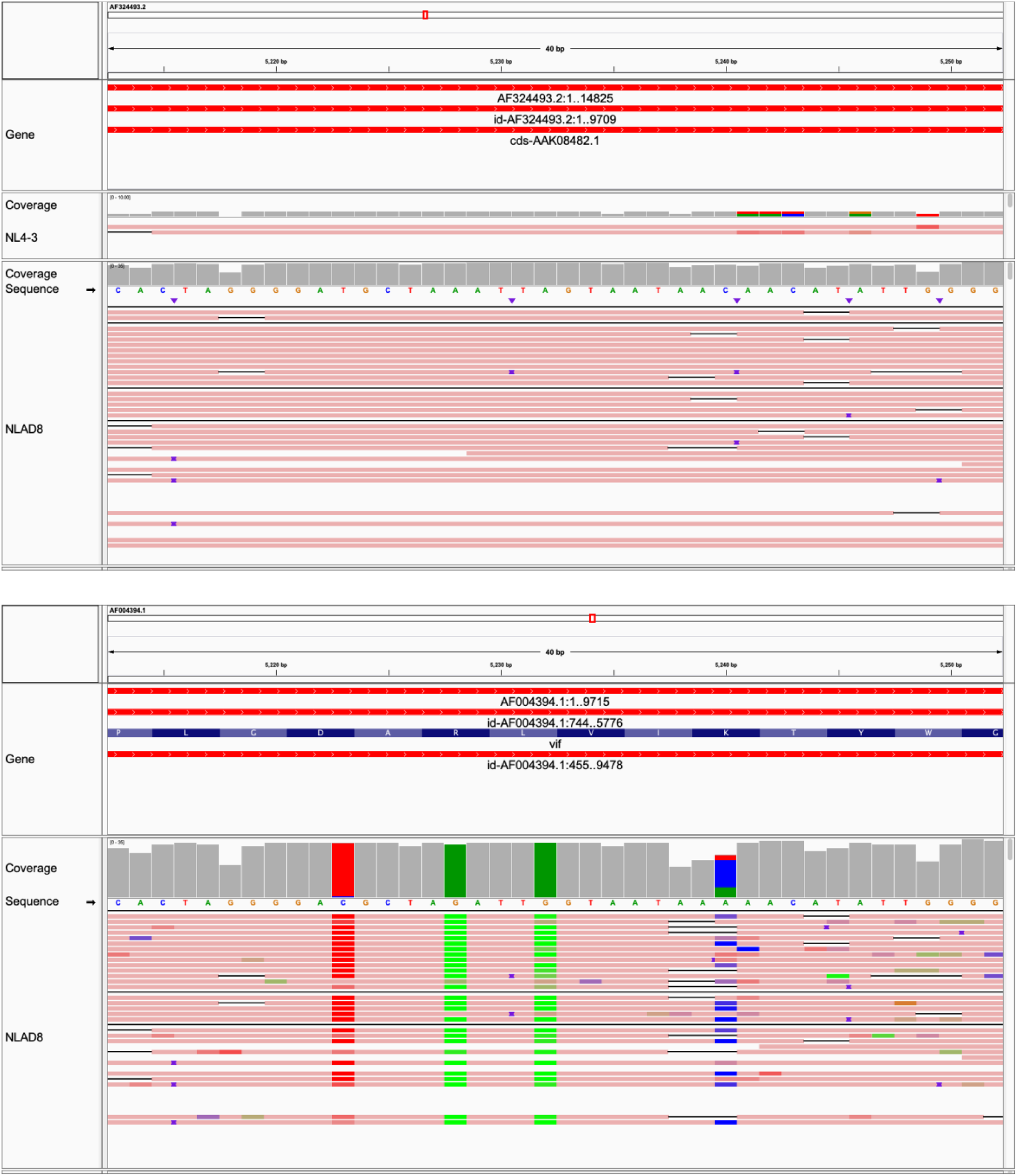
SNVs near an A-homopolymer truncation. Top: AF324493.2:5,213-5,252. This region is common between NLAD8 and NL4-3 (top), and we expect differences in parts of the virus outside the *env* gene (bottom). In above, we see 4 SNVs, one embedded in a homopolymer track. The fact that we can read across such a large A-homopolymer (thus more easily discounting the false positive deletions in reads at the A-homopolymer) may either be due to a SNV breaking up the A-homopolymer, or less likely a shift in the relative position of C. Compare to SAME Region in NL4-3. Bottom: AF004394.1:5,213-5,252. Note that the usage of SNVs here is to denote differences between reference samples NLAD8 and AD8, and not SNVs in NLAD8. Alignments with minimap2 [13] in usegalaxy.eu [14]. Visualized in Integrative Genomics Viewer [49].

**Supplemental Figure 4:**
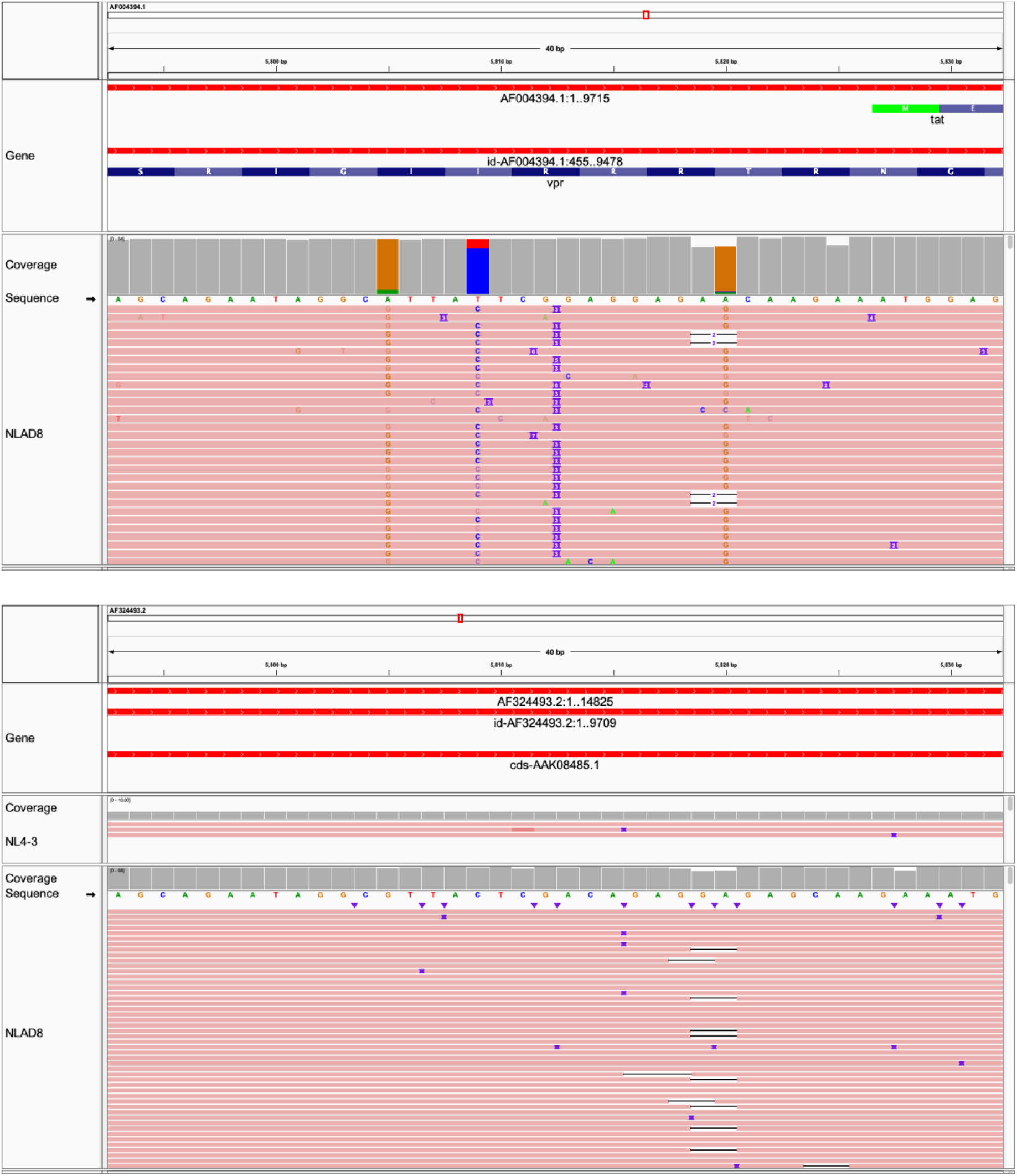
Insertion. Top: AF004394.1:5,793-5,832. This region is common between NLAD8 and NL4-3, and we expect differences in parts of the virus outside the *env* gene. Compare to NL4-3. Bottom: AF324493.2:5,793-5,832. Alignments with minimap2 [13] in usegalaxy.eu [14]. Visualized in Integrative Genomics Viewer [49].

**Supplemental Figure 5:**
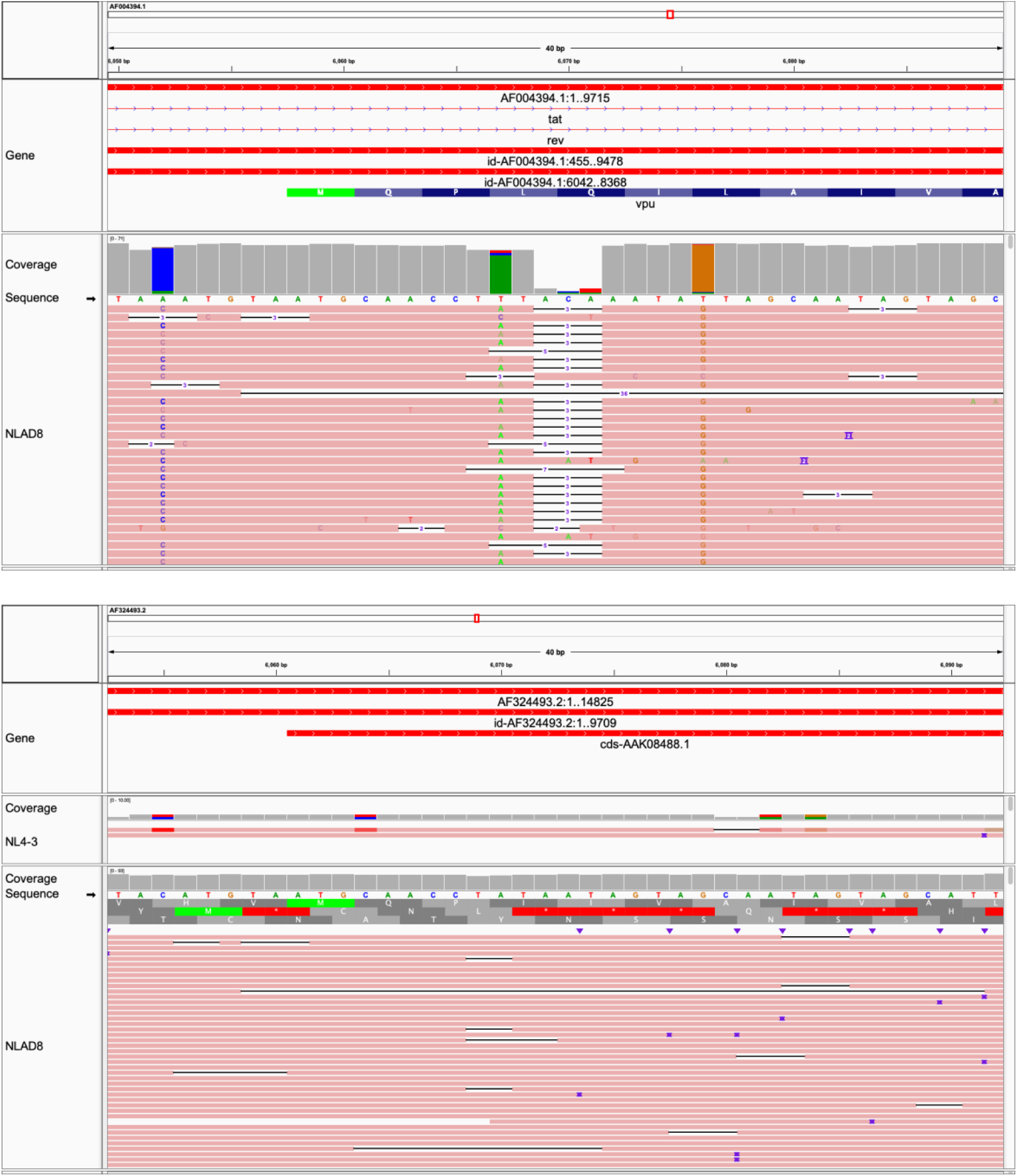
A likely true positive deletion with respect to reference AD8 in NLAD8. This region is common between NLAD8 and NL4-3, and we expect differences in parts of the virus outside the *env* gene. AF004394.1:6,050-6,089. While technically out of frame, given the context of the above, this deletion would cause an in-frame substitution of LQ at amino acids 4 and 5 and gain of I at position 4 in vpu. Compare to NL4-3. Alignments with minimap2 [13] in usegalaxy.eu [14].Visualized in Integrative Genomics Viewer [49].

**Supplemental Figure 6:**
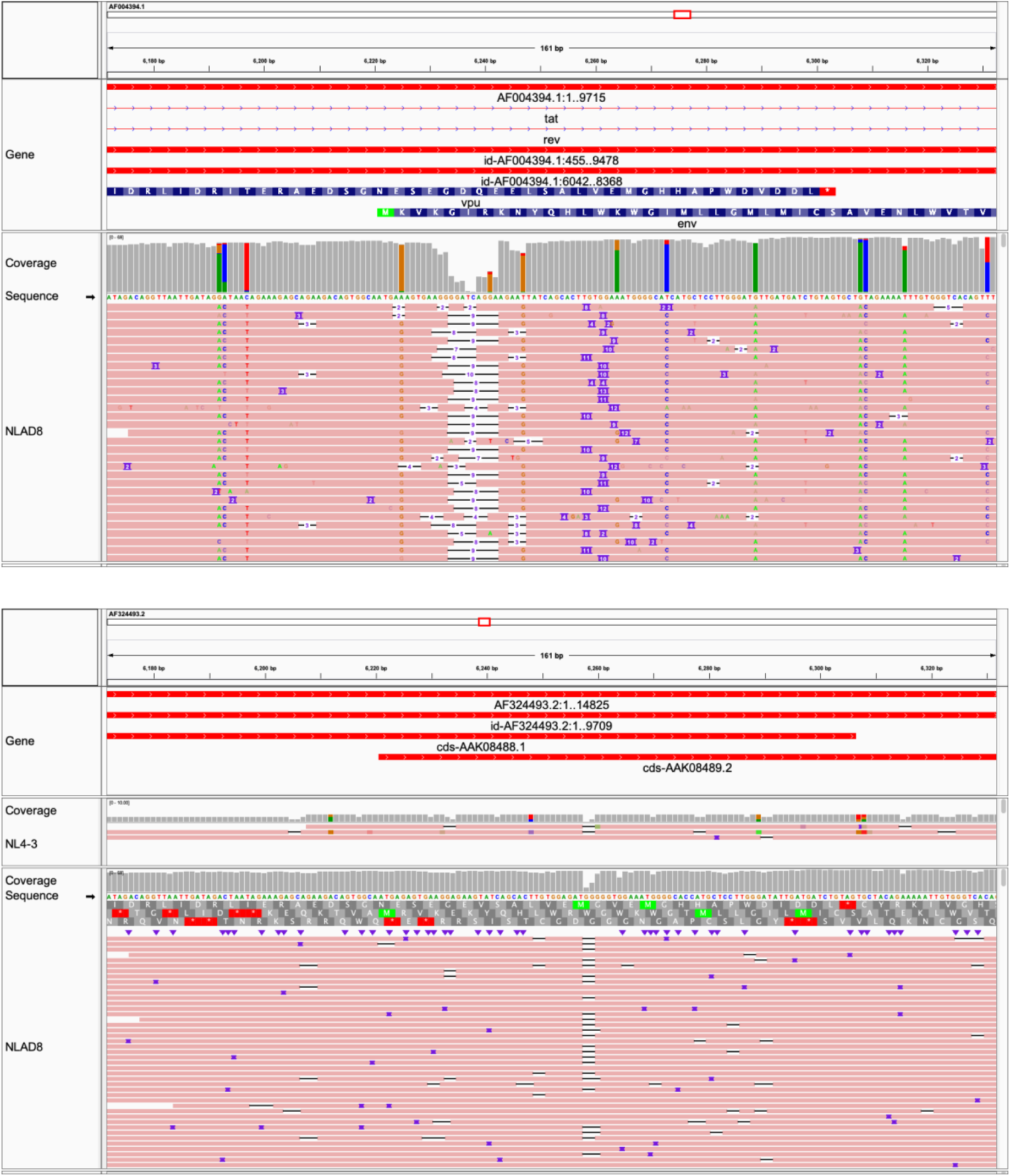
Structural variants intermingled with SNVs. Top: AF004394.1:6,172-6,332. Bottom: AF324493.2:6,172-6,332. Region conserved between NL4-3 and NLAD8 (bottom), but not between NLAD8 and AD8 (top). Alignments with minimap2 [13] in usegalaxy.eu [14]. Visualized in Integrative Genomics Viewer [49].

**Supplemental Figure 7:**
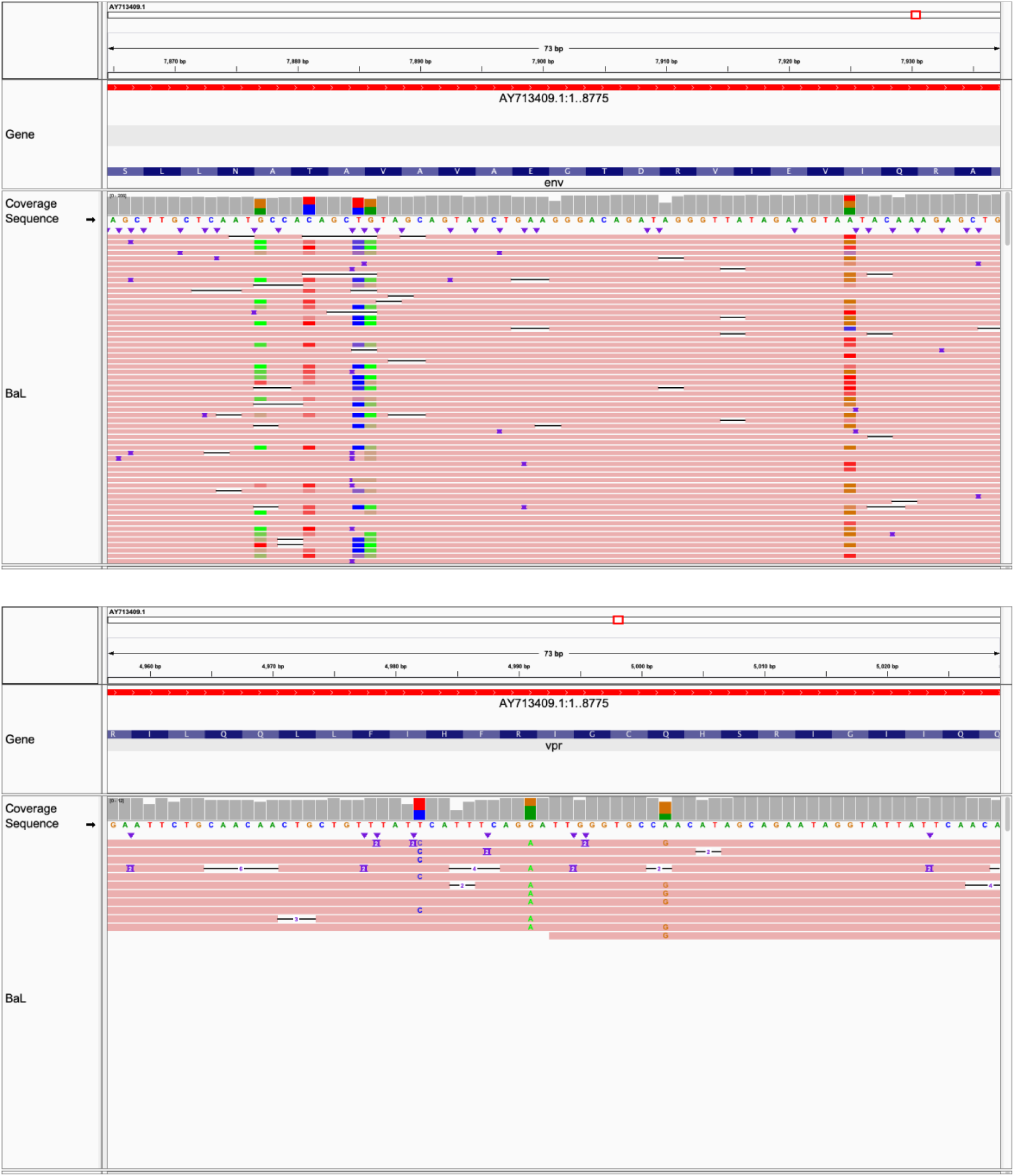
SNVs define HIV haplotypes directly. Haplotypes in BaL are observed to be mutually exclusive. Top: AY713409.1:7,865-7,937. Bottom: AY713409.1:4,957-5,029. Note two haplotypes are seen here. One as g.4982T>C, another as g.4991G>A with g.5002A>G (other phased SNVs not mentioned). Alignments with minimap2 [13] in usegalaxy.eu [14]. Visualized in Integrative Genomics Viewer [49].

**Supplemental Table 3:**
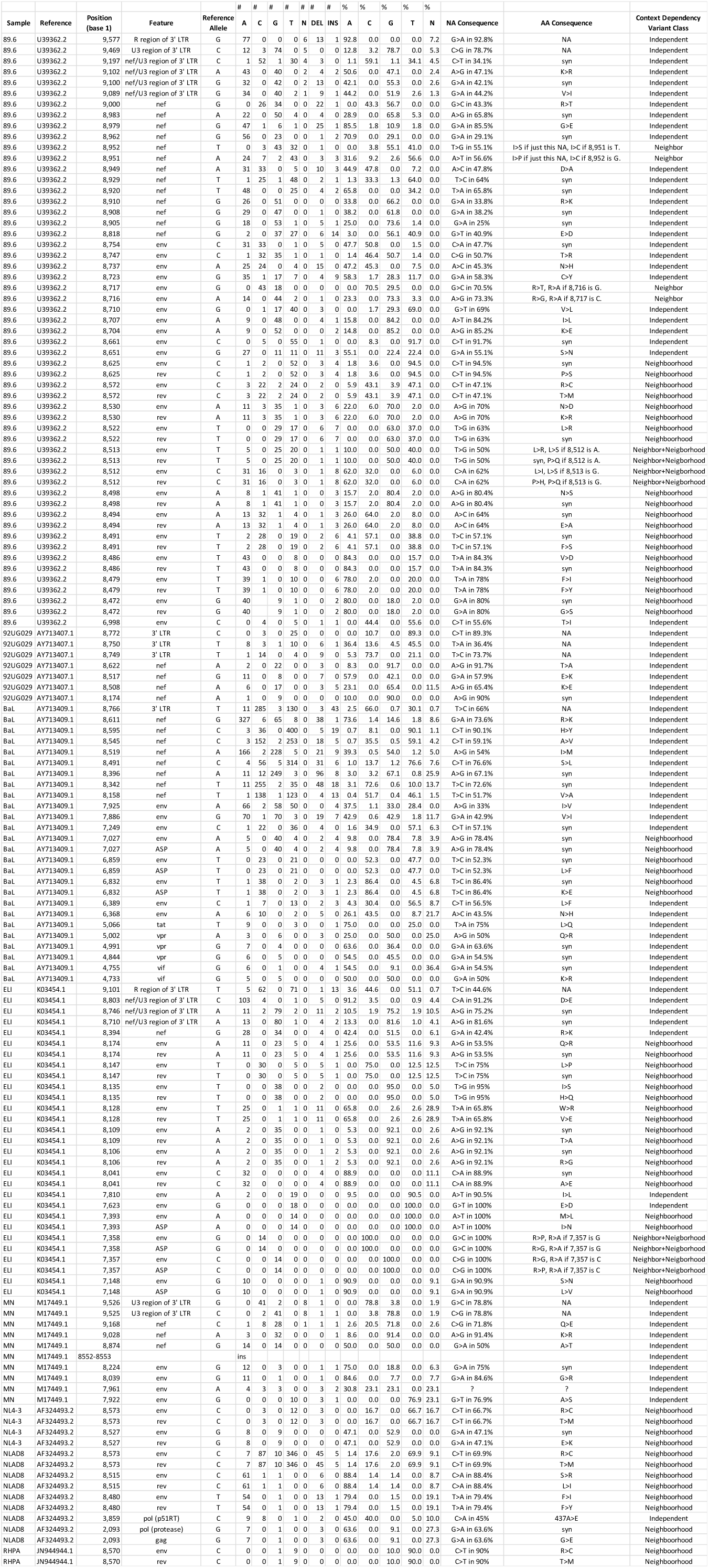
SNVs in reference HIV-1 strains. NA: Nucleic Acid. AA: Amino acid. ASP: Antisense protein. LTR: Long terminal repeat. RT: Reverse transcriptase.

**Supplemental figure 8:**
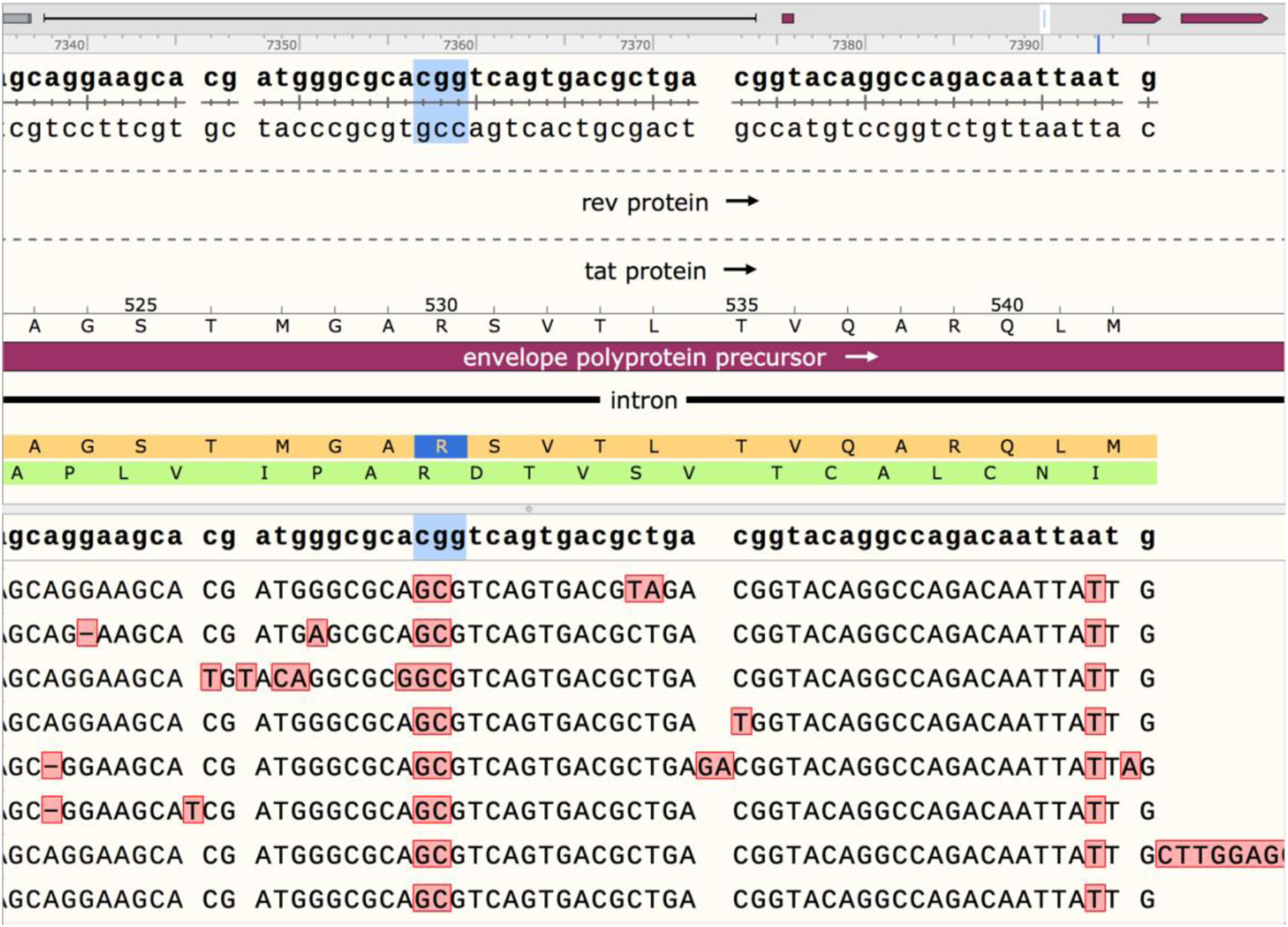
Context Dependency Variant Classification. Independent = a variant whose effect is limited to its coding frame. Neighbor = a variant whose effect is limited to its coding frame, but when considering variation in its adjacent nucleotide, can contribute to a codon change dependent on both states. Neighborhood = a variant whose effect is not limited to its coding frame (example: when proteins are coded by the same mRNA but in different frames as part of *gagpol*, *tat*/*rev*/*env*/*vpu*/ASP), but when considering its adjacent nucleotide, can contribute to a pair of codons changing dependent on both states. Neighbor+Neighborhood = when Neighbor and Neighborhood dependencies are satisfied. Here we have a frame from ELI, where Neighbor+Neighborhood occurs at R530 (highlighted in blue) in env (maroon and orange in forward orientation), and also overlaps residues in Antisense protein (in green, negative orientation).

**Supplemental Figure 9:**
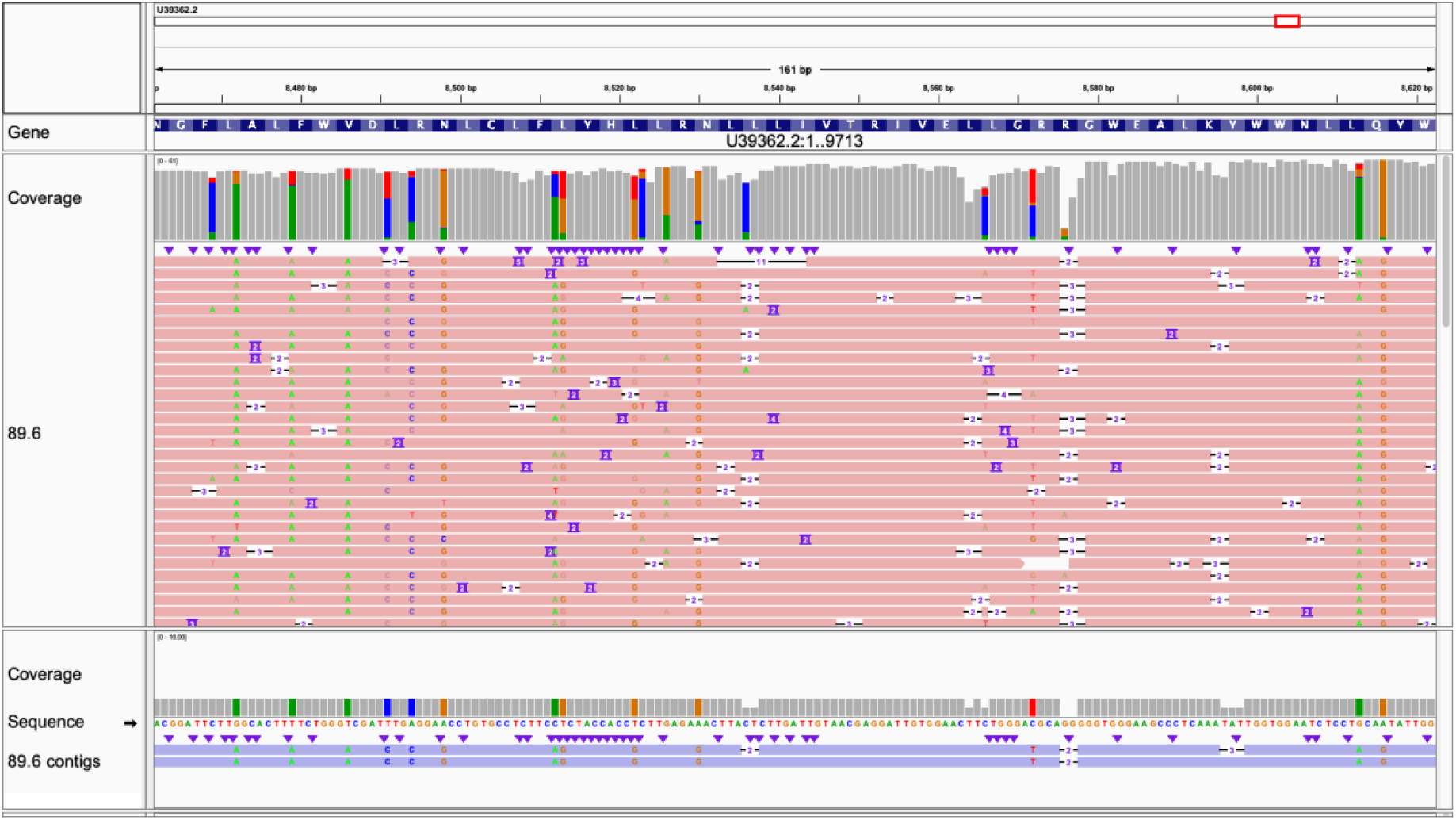
Canu *de novo* assembly supports manual SNV calling. 89.6. U39362.2:8,462-8,622. Note that contigs were assembled in negative orientation. Alignments with minimap2 [13] in usegalaxy.eu [14]. Visualized in Integrative Genomics Viewer [49].

**Supplemental Figure 10:**
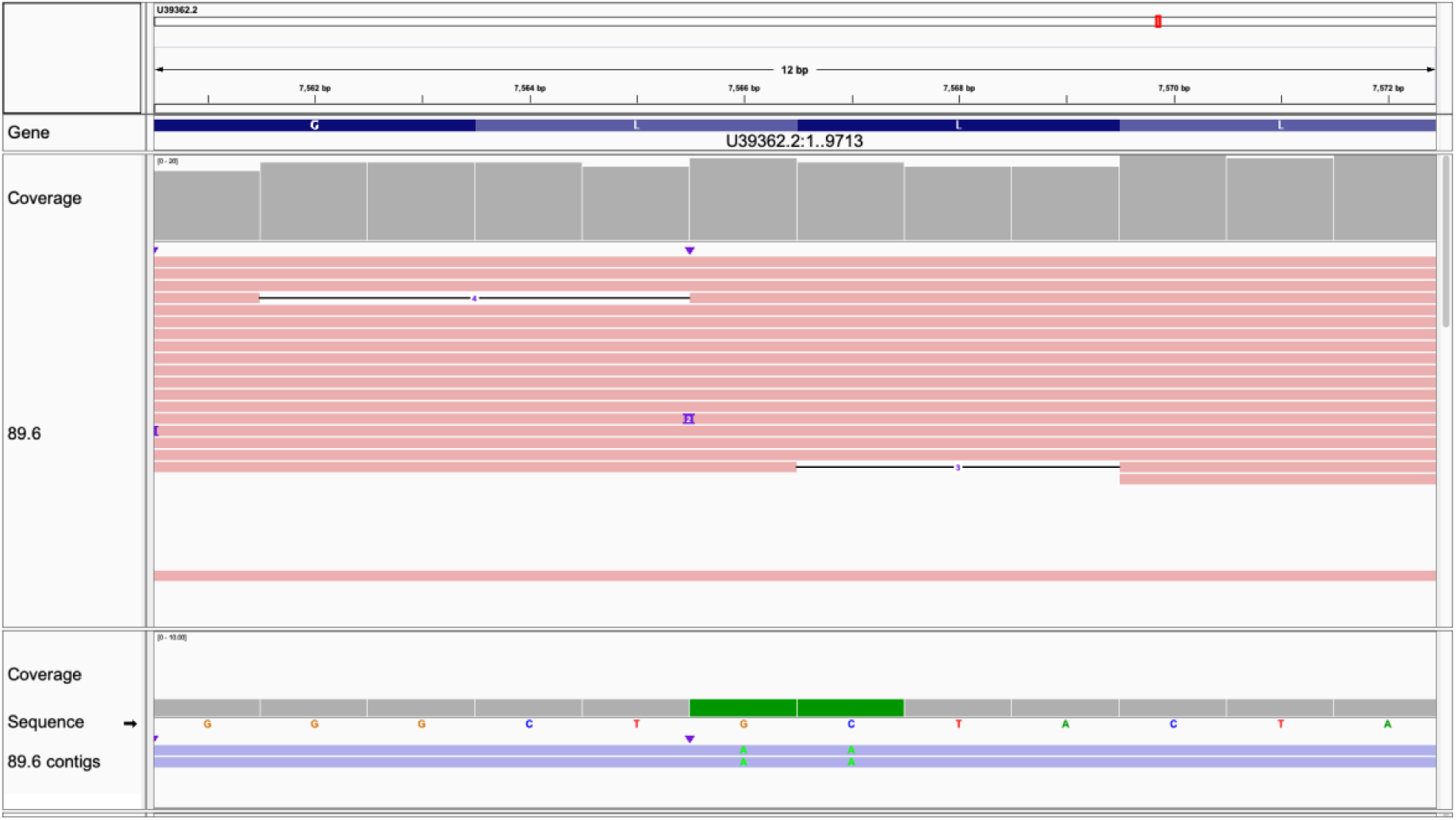
Canu *de novo* assembly can incompletely resolve haplotypes. 89.6. U39362.2:7,561-7,572. SNVs in assemblies not supported by visual inspection. Left G: 4/14 A. Right C: 4/14 A. Note that contigs were assembled in negative orientation. Alignments with minimap2 [13] in usegalaxy.eu [14]. Visualized in Integrative Genomics Viewer [49].

**Supplemental Figure 11:**
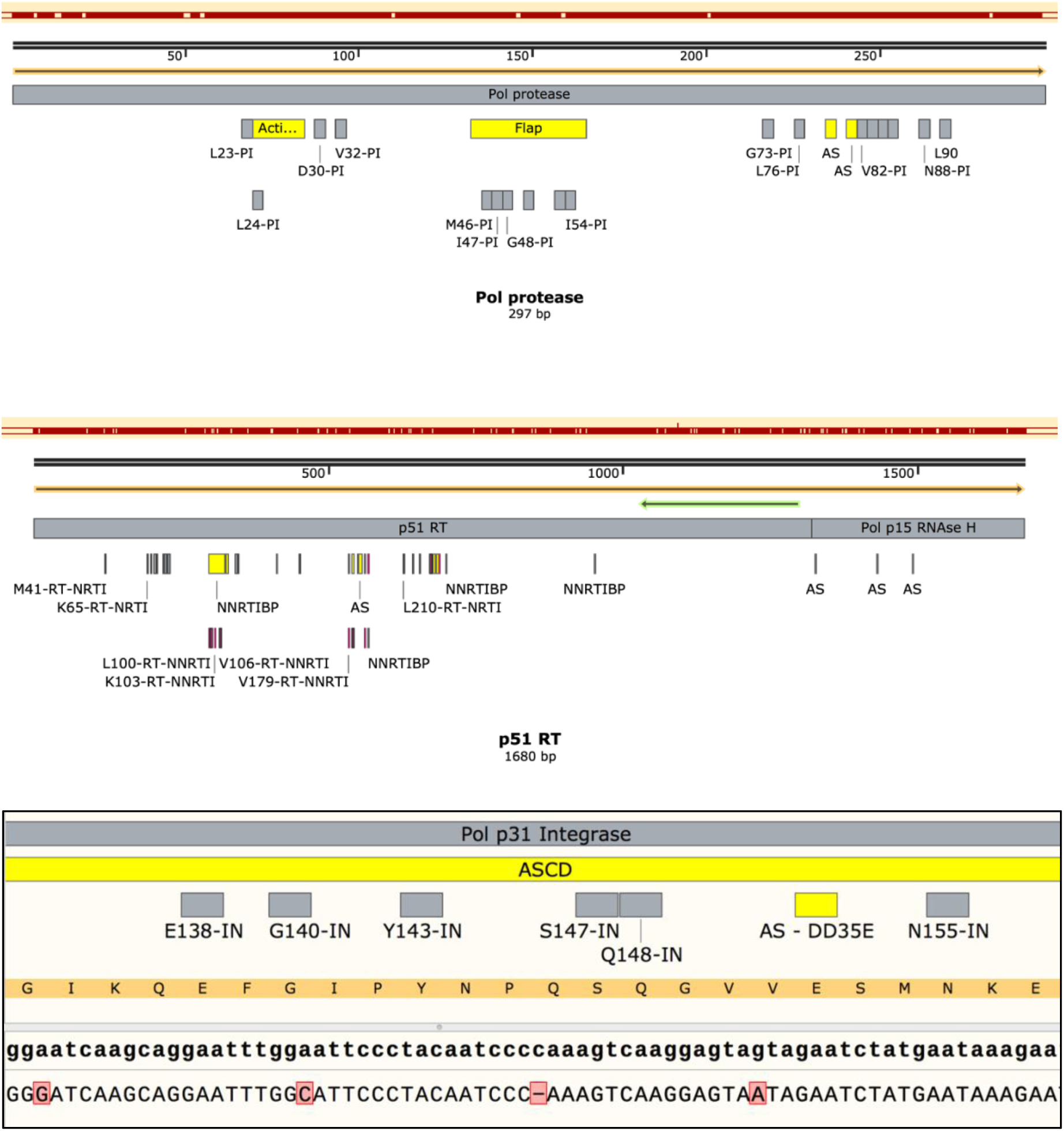
*De novo* contigs can provide drug resistance information. Drug resistance data from Stanford HIV Drug Resistance Database [23, 24]. Reference HXB2 K03455.1. Maroon line is the mapped NLAD8 contig. Close-up of integrase shows silent mutation at G140G and a missense mutation at V151I, both around known resistance-associated residues. The truncated C homopolymer in the center of the viewing window is most likely an ONT artifact. Visualized in SnapGene version 5.0.4.

**Supplemental Figure 12.**
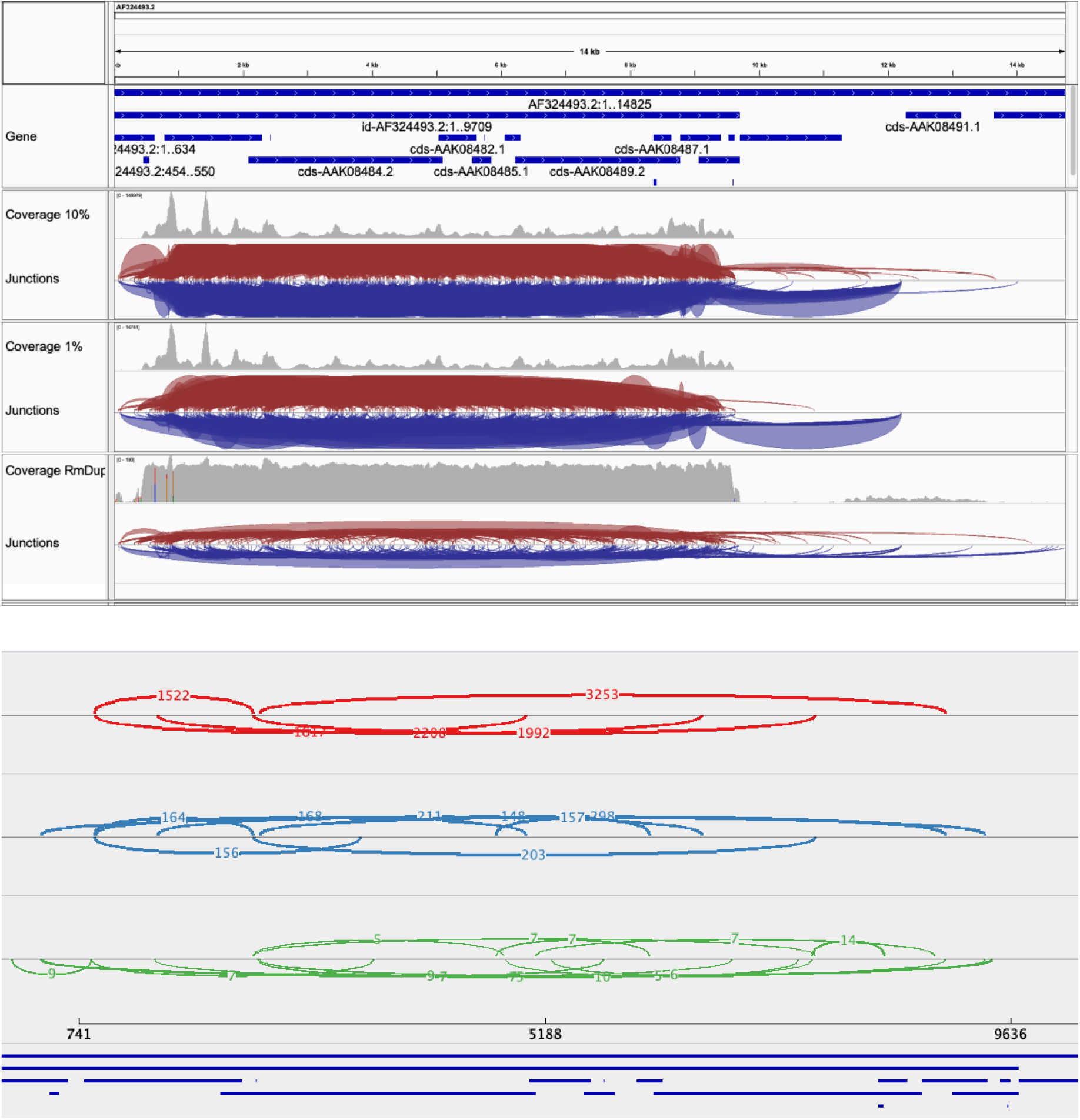
Short-read NL4-3 virion RNA-seq. Single-end 76 bp. SRR3472915. Reference: pNL4-3 AF324493.2. Note backbone contamination, either due to backbone DNA contamination or backbone insertion and expression. Forensic genomic analyses support cDNA + PCR library, although this information was not available from SRA entry. Top: mapped with HISAT2 (a split-read aligner/mapper [50]). Note abundance of forward and reverse splice junctions. Bottom: Sashimi plot of forward splice junctions only. Red: data downsampled to 10%. Minimum splice junction frequency set to 1470. Blue: data downsampled to 1%. Minimum splice junction frequency set to 147. Green: data processed with RmDup to remove PCR duplicates. Minimum splice junction frequency set to 5. Most reads include 3’LTR elements, and a canonical exon upstream of *gag*.

## LIST OF DATA (To be deposited in SRA or ENA in near future)

Basecalled files

.dna files of contigs (n=?)

AF033819.3-anchored .bam/.bai files

Strain-specific .bam/.bai files

Any multi-sequence alignments

## Notes

#### Summary of Updates

A citation error between [1] and [3] was corrected. A citation [21] referencing "camouflaged" regions was added to attribute the terminology describing the concept to Ebbert et al. 2019. Minor typos were addressed. Author affiliations were reformatted.

